# The m^6^A reader IGF2BP2 directs immune-metabolic reprogramming in Leishmania amazonensis-infected macrophages

**DOI:** 10.1101/2022.09.08.507100

**Authors:** Sheng Zhang, Hervé Lecoeur, Hugo Varet, Rachel Legendre, Nassim Mahtal, Caroline Proux, Nathalie Aulner, Spencer Shorte, Capucine Granjean, Philippe Bousso, Eric Prina, Gerald F. Späth

## Abstract

Macrophages are the major host cells of the protozoan parasite *Leishmania* in mammalian infection. These key innate immune cells display remarkable phenotypic plasticity ranging from pro-inflammatory M1 to anti-inflammatory M2 macrophages that can control infection and tissue homeostasis, respectively. It has been recognized that *Leishmania* exploits macrophage phenotypic plasticity to establish chronic infection. However, the current notion that these parasites simply trigger an M2-like phenotype seems over-simplified considering the immunopathology observed during leishmaniasis – in particular in response to *Leishmania amazonensis* - which is often characterized by a mixed Th1/Th2 immune response. Here we combined a series of systems-level analyses to shed new light on the phenotype of *Leishmania*-infected macrophages (LIMs) during short- and long-term infection, *in vitro* and *in vivo*. Immuno-metabolic profiling by RNA-seq, RT-qPCR, cytokine immunoassays, and real-time bioenergetic flux analysis of *L. amazonensis*-infected bone marrow-derived macrophages (BMDMs) revealed a highly complex and unique phenotypic and bioenergetic signature. *In vitro* LIMs were characterized by co-expression of both M1 and M2 markers at RNA and protein levels and increased expression of glycolytic genes that matched a progressive metabolic switch from a M2-like respiratory to a M1-like glycolytic energy production observed for both long-term *in vitro* and *in vivo* infected macrophages. Unlike in M1 macrophages, glycolytic gene expression did not correlate with increased expression of its key regulatory HIF-1α. In contrast, siRNA knock down experiments in primary BMDMs uncovered an essential role of the m^6^A reader protein IGF2BP2 in stabilizing m^6^A modified transcripts of the glycolytic pathway, contributing to HIF-1α-independent induction of glycolysis. In conclusion, *L. amazonensis* establishes a complex and unique phenotypic shift in infected macrophages *in vitro* and *in vivo* that combines M1-like and M2-like immuno-metabolomic characteristics and implicates differential mRNA stability in induction of aerobic glycolysis. Our data thus uncover epi-transcriptomic regulation as a novel target for *Leishmania* immune subversion to establish a host cell phenotype beneficial for intracellular parasite development and chronic infection.

## INTRODUCTION

Intracellular pathogens have co-evolved strategies to neutralize anti-microbial activities of their host cells, harness their nutritional supplies and domesticate their metabolism to promote microbial survival and proliferation. The eukaryotic pathogen *Leishmania* represents an interesting model system to investigate such immune-metabolic subversion strategies as these parasites infect and thrive inside important immune cells, including macrophages (Mφs) (Antonia and Ko, 2020; Bogdan, 2020; Lecoeur et al., 2022). These key innate immune cells display remarkable phenotypic plasticity that adapts their immune functions to local tissues and changes in the microenvironment. Indeed, Mφs can adopt a spectrum of polarization states ranging from canonical pro-inflammatory M1 cells that carry antimicrobial immune function and promote T helper type 1 responses, to anti-inflammatory M2 cells that promote T helper type 2 responses and are associated with tissue homeostasis and wound-healing. Mφ polarization is driven by a complex interplay between extrinsic immune signals (microbial agents, cytokines/chemokines) and intrinsic metabolic signals. On one hand, M1 polarization is triggered by pro-inflammatory cytokines and linked to induction of (i) HIF-1α-dependent glycolytic activity that rapidly generates ATP (Corcoran and O’Neill, 2016), (ii) the Pentose Phosphate Pathway (PPP), (iii) fatty acid synthesis, and ii) an interrupted Tricarboxylic Acid (TCA) cycle (Russell et al., 2019). On the other hand, M2 polarization is triggered by anti-inflammatory cytokines and is associated with (i) an intact TCA cycle, (ii) increased mitochondrial oxidative phosphorylation (OXPHOS), (iii) increased fatty acid oxidation, and (iv) the suppression of the PPP (Diskin and Palsson-McDermott, 2018).

The capacity of *Leishmania* to subvert Mφ immune functions has attracted considerable attention given its important role in causing the severe immuno-pathologies characteristic of leishmaniasis. Considering various parasite species, hosts, Mφ systems and infection methods, the prevailing model suggests that *Leishmania* triggers M2 polarization to establish permissive conditions for infection through the reduced expression of pro-inflammatory genes (e.g. *Nos2*, inflammasome components) and increased expression of key molecules such as arginase-1 (Kumar et al., 2018; Lecoeur et al., 2020). In contrast, the interaction of *Leishmania* with the host cell metabolisms and the impact of metabolic reprogramming on Mφ polarization is less investigated, even though M1 and M2 states have tailored metabolic profiles that support divergent macrophage functions (Cohen and Mosser, 2013; Murray, 2017; O’Neill et al., 2016; Russell et al., 2019). *Leishmania* were shown to subvert various Mφ metabolic pathways during short-term infection *in vitro*, switching from an early glycolytic metabolism to oxidative phosphorylation (Moreira et al., 2015; Rabhi et al., 2012; Ty et al., 2019), modulating cholesterol and triglyceride homeostasis (Rabhi et al., 2012), and causing accumulation of lipid droplets (Rodriguez et al., 2017). However, no detailed information is available on how *Leishmania* fine-tunes the interplay between Mφ polarization state, immuno-expression, and metabolic functions. Furthermore, current studies rely exclusively on short-term, *in vitro* infection models using the promastigote developmental stage, all of which only poorly mimics the immune-metabolomic state of chronically infected Mφ *in vivo* harboring hundreds of intracellular amastigotes (Lecoeur et al., 2020).

To overcome these limitations, we conducted an integrative, immuno-metabolomic analysis *in vitro* using murine bone marrow-derived macrophages (BMDMs) infected with lesion-derived *L. amazonensis* amastigotes for up to 30 days post-infection, and *in vivo* using infected tissue Mφs isolated from lesions for *ex vivo* studies. Integration of transcriptomic, bioenergetic and proteomic profiles uncovered a previously under-appreciated, highly complex phenotype of *Leishmania*-infected macrophages (here termed LIMs) far from a stereotypic M2-like polarization state. We provide strong evidence that chronically infected LIMs co-express M1 and M2 immuno-metabolic features that may be at the origin of the equally complex T cell response observed during clinical *Leishmania* infection. We further identify the epi-transcriptomic reader protein IGF2BP2 as an important component in HIF-1α−independent induction of aerobic glycolysis in LIMs that relies on stabilization of *Hk2* mRNA by increased m^6^A methylation.

## MATERIAL AND METHODS

### Ethics statement

Six-week-old female C57BL/6 and Swiss *nu/nu* mice were purchased from Janvier (Saint Germain-sur-l’Arbresle, France). Rag2^tm1Fwa^ (Shinkai et al., 1992) were bred at the Institut Pasteur. All animals were housed in A3 animal facilities according to the guidelines of Institut Pasteur and the “Comité d’Ethique pour l’Expérimentation Animale” (CEEA) and protocols were approved by the “Ministère de l’Enseignement Supérieur; Direction Générale pour la Recherche et l’Innovation” under number #19683.

### Isolation, culture, and polarization of bone marrow-derived macrophages

Bone marrow cells were recovered from tibias and femurs of C57BL/6 mice and cultured in complete medium supplemented with recombinant mouse Colony Stimulating Factor 1 (rm-CSF-1) to give Bone Marrow-Derived Macrophages (BMDMs) as previously described (Lecoeur et al., 2020) (Supplementary Material and Methods). After 6 days adherent Mφs were detached with 25 mM EDTA in PBS, seeded in complete medium and treated during three days with different cytokine cocktails to induce specific polarization states: i) 30 ng/mL rm-CSF-1 for M0, ii) sequential stimulation with 50 ng/mL LPS (*E. coli* strain 0111: B4; Alpha Diagnostic, 11-1) for 2 days and with 20 ng/mL IFN-γ (ImmunoTools, 681588) for 1 day for classically activated (M1) BMDMs, and iii) 50 ng/mL rm-CSF-1 plus 10 ng/mL IL-4 (ImmunoTools, 681680) and 10 ng/mL IL-13 (ImmunoTools, 681300) for 3 days for alternatively activated (M2) BMDMs.

### Isolation of lesional macrophages from mouse footpads

Lesional Mφs were isolated from footpad lesions of RAG2 knock-out mice previously inoculated with lesion-derived amastigotes. These *in situ* infected macrophages were used for *ex vivo* analyses (and therefore referred to as *ev*LIMs). Mice were euthanized by carbon dioxide inhalation, footpads were removed and placed in Digestion Buffer (DB) composed of 50 U/ml DNAse I (Sigma-Aldrich), 100 U/ml collagenase II (Sigma-Aldrich), 100 U/ml collagenase IV (Sigma-Aldrich) and 1 U/ml dispase II (Roche Applied Science) in DMEM and placed on a 100 µm cell strainer. Footpads were perfused with one ml of DB at different locations and maintained at 37°C for 30 minutes. This step was repeated two times. Macrophages were recovered and washed in PBS (centrifugation at 50 g, 10 min, 4°C) before processing.

### *Leishmania* isolation and BMDM infection

mCherry transgenic, tissue-derived amastigotes of *Leishmania amazonensis* strain LV79 (WHO reference number MPRO/BR/72/M1841, *L. am*) were isolated from infected footpads of Swiss nude mice. To establish *in vitro* infected macrophages (*iv*LIMs), an MOI of four amastigotes per host cell was used. *iv*LIMs were cultured at 34°C as previously described and subjected to various analyses at 3, 15 and 30 days post-infection to assess short-, mid- and long-term infections, respectively (Lecoeur et al., 2010).

### Determination of the bioenergetic profile of macrophages and purified amastigotes

Bioenergetic profiles for M0, M1, M2 Mφs, *iv*LIMs, and *ev*LIMs were established by assessing glycolytic-*vs* OXPHOS- (mitochondrial oxidative phosphorylation) dependent ATP production using the Agilent Seahorse XF Real-Time ATP Rate Assay Kit (Agilent, 103592-100). The mitochondrial function of macrophages and amastigotes was analyzed by the Agilent Seahorse XF Cell Mito Stress Test Kit (Agilent, 103015-100). The glycolytic function of infected macrophage was analyzed by the Agilent Seahorse XF Glycolytic Stress Test Kit (Agilent, 103020-100). Mφs and amastigotes were distributed in Seahorse XF96 tissue culture microplates for 3 days at 34°C. For experiments with free parasites, lesion-derived amastigotes were seeded onto poly-L-lysine (0.1 µg/mL, 350 kDa) coated wells prior to analysis by the Seahorse XFe96 analyzer according to the manufacturer’s conditions. Modifications of the protocols and consequent analyses are described in Supplementary Material and Methods.

### Quantification of nitrites

Nitrite quantitation in culture supernatants was performed by the colorimetric Griess test kit (Thermo Fisher Scientific, G7921) according to the manufacturer’s instructions (Supplementary Material and Methods).

### Quantitation of extracellular lactate

Lactate levels of supernatants derived from M0, M1, M2, *iv*LIMs and *ev*LIMs cultures, were quantified using the L-lactate Assay (Sigma-Aldrich, MAK064) and D-lactic acid colorimetric Assay (elabscience, E-BC-K002-M) kits in accordance with the manufacturer’s instructions. Results were normalized for cell density.

### Sample preparation for Western Blotting (WB)

Cell or amastigote samples were homogenized in ice-cold CelLytic^TM^ M buffer (Sigma-Aldrich, C2978) with 1x protease inhibitor cocktail (ThermoFisher Scientific, 78442). After a centrifugation at 14,000 g for 10 minutes, supernatants were collected, and protein concentration was determined by the BCA protein assay kit (Abcam, ab287853). Then samples were aliquoted and stored at −80°C. Western blot analyses were performed as described in Supplementary Material and Methods.

### Sample preparation for the Proteome Profiler Mouse XL Cytokine Array

Supernatants of Mφ cultures from 3 independent biological replicates were collected, pooled, and analyzed by the Proteome Profiler Mouse XL Cytokine Array (ARY028, Biotech, R&D systems) according to the manufacturer’s instructions. Revelation and analysis were performed as described in Supplementary Material and Methods.

### Morphologic analysis of BMDM cultures

Infected and differentially polarized BMDMs initially seeded in 384-well plates (CellCarrier™, Perkin Elmer) were analyzed using the confocal OPERA Phenix® system. On day 3 post-infection or post-polarization, BMDMs were fixed and analyzed according to protocols described in Supplementary Material and Methods.

### Sample preparation for RT-qPCR analysis

Total RNA isolation, quality control, and reverse transcription were performed using the NucleoSpin® RNA plus kit (Macherey-Nagel, 740984.50) following the manufacturer’s instructions. Polarized BMDMs and *iv*LIMs were cultured in 6-well plates (Corning Life Science, 353046) and were homogenized in lysis buffer. *ev*LIMs were pelleted right after isolation and resuspended directly in the lysis buffer. Then samples were treated and analyzed as detailed in Supplementary Material and Methods. The list of primers used for the RT-qPCR analyses is provided in Supplementary Table 3.

### Sample preparation for RNA-seq analysis

RNA isolation was performed with the RNeasy^+^ isolation kit (Qiagen) according to the manufacturer’s instructions. Evaluation of RNA quality was carried out by optical density measurement using a NanoDrop spectrophotometer (Kisker) as previously described (Lecoeur et al., 2020). Total RNA was isolated from three independent biological replicates of M0 BMDMs and *iv*LIMs at day 3 post-infection and treated as described in Supplementary Material and Methods.

### *In silico* analyses

The ClusterProfiler package (version 4.2.2) of R was used for gene set enrichment analysis (GSEA) and for data mapping on metabolic pathways using the Kyoto encyclopedia of genes and genomes (KEGG) database. The gene set variation analysis (GSVA) package (version 1.48.3) of R was used to assess GO-term enrichment in our data sets. Results were visualized with the enrichplot (version 1.14.1), pathview (version 1.34.0) and ggplot2 (version 3.4.4) packages. The public database POSTAR3 was used to explore RNA Binding Protein (RBP) – mRNA interactions (Zhao et al., 2022) in the context of our RNA-seq experiment. Protein-RNA interaction networks were visualized in Cytoscape (version 3.9.1). A list of M1 and M2 markers was compiled and manually curated from published RNA-seq studies (Dichtl et al., 2021; Jablonski et al., 2015; Orecchioni et al., 2019; Zhang et al., 2019) (Supplementary Tables 1 and 2).

### Gene knock-down by small RNA interfering (siRNA)

Knock down of *Igf2bp2* transcripts was performed with two specific siRNAs (siRNA-1, GACUACUCCUUCAAGAUUU; siRNA-2, CAUCUCAUCCUUGCAGGAU). Universal scrambled siRNA duplexes were used as negative control (SIC001) (Sigma-Aldrich, Germany). siRNAs were transfected into M0 macrophages or *iv*LIMs using the GenMuteTM siRNA Transfection reagent (SignaGen Laboratories, SL100568-PMG) as specified by the manufacturer. Differentiated BMDMs were seeded in 6-well plates (Corning Life Science, 353046, 3×10^6^ per well), infected or not with *L. am* and cultured at 34°C for 24 hours until transfection (Lecoeur et al., 2020). The medium of each well was replaced with 1 mL of fresh medium 60 minutes before transfection. 100 pmoles siRNA were mixed with 100 μL 1X transfection buffer and incubated with 4 μL transfection reagent for 15 minutes at room temperature, under gentle rocking. After 3 hours of incubation at 34°C, the transfection medium was replaced with fresh, pre-warmed complete medium. Transfection efficiency of siRNA for *Igf2bp2* knock down studies was tested on M0 macrophages by RT-qPCR and Western blot analysis.

### RNA immunoprecipitation (RIP) and Methylated (m^6^A) RNA immunoprecipitation (MeRIP)

RNA immunoprecipitation and Methylated (m^6^A) RNA immunoprecipitation were performed in M0 or *iv*LIMs according to modifications of initial protocols (Dominissini et al., 2013; Gagliardi and Matarazzo, 2016) as detailed in Supplementary Material and Methods.

### Statistical analyses and graphical representations

Statistics were determined by the t-test or one-way ANOVA using the GraphPad Prism 10.2.1 software. Graphs were generated using R 4.3.2, GraphPad Prism 10.2.1 or Cytoscape 3.10.0.

## RESULTS

### *In vitro Leishmania*-infected macrophages (*iv*LIMs) exhibit a distinctive immune expression pattern characterized by a combination of M1, M2, and LIMs-specific transcriptomic features

*In vitro* LIMs (*iv*LIMs) were obtained by infecting C57BL/6 bone marrow-derived macrophages (BMDMs) with virulent, lesion-derived *Leishmania amazonensis* amastigotes (Lecoeur et al., 2020). Following 3 days of infection, *iv*LIMs were compared to M0 (unpolarized) macrophages at day 3 post-differentiation, as well as to M1 and M2 macrophages at day 3 post-polarization generated using established culture conditions (Supplementary Figure 1A and B). *iv*LIMs exhibited a distinct morphology characterized by a highly irregular cell shape and the presence of large parasitophorous vacuoles containing parasites (Figure 1A). *iv*LIMs neither resemble M1 macrophages, which adopted a typical amoeboid shape (Figure 1A), harbored lipid droplets and produced nitrites (Supplementary Figure 1C and D), nor M0 or M2 macrophages, which exhibited an elongated cell shape (Figure 1A).

**FIGURE 1.**
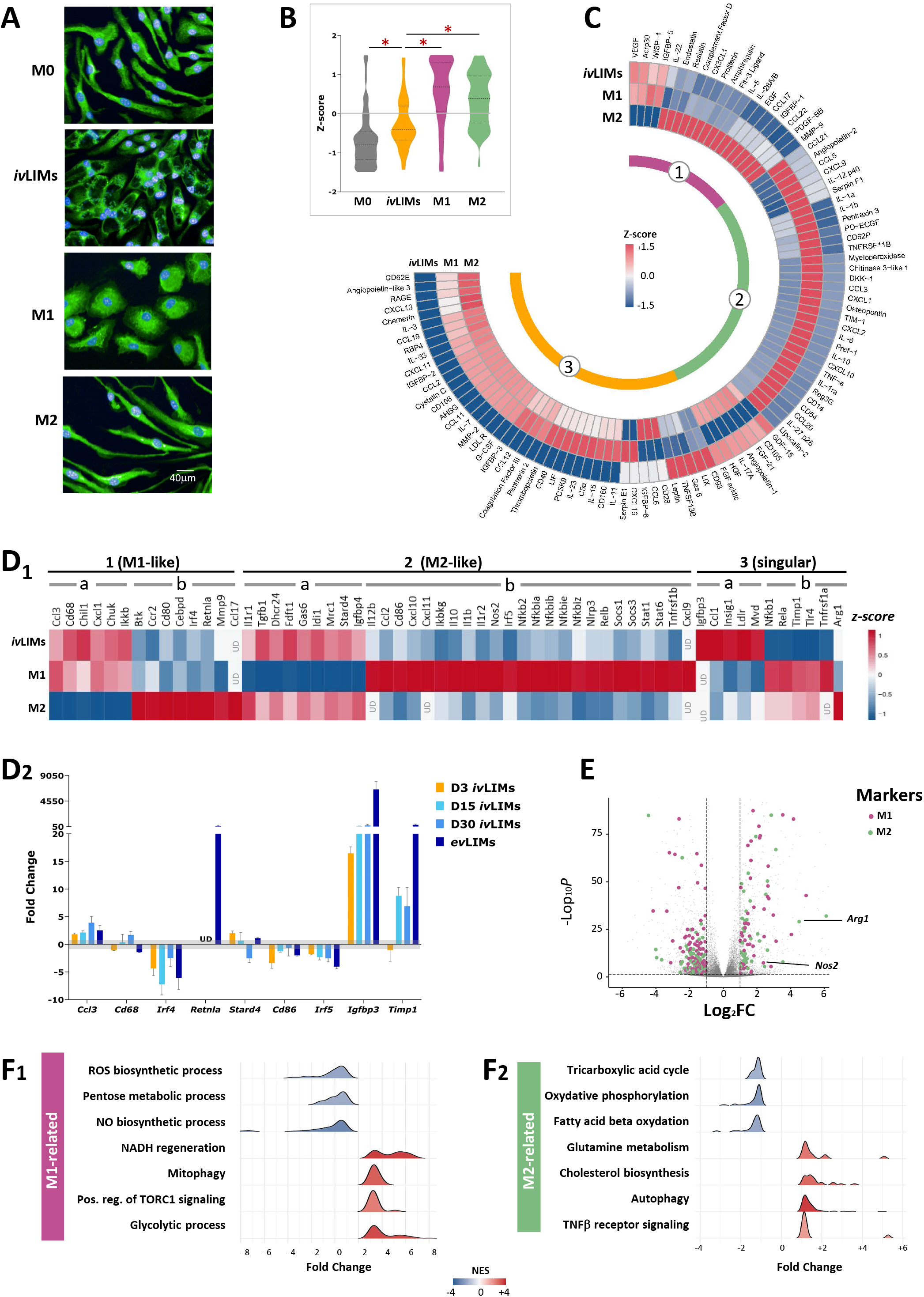
Phenotypic characterization of *L. amazonensis-*infected macrophages. **A)** Analysis of macrophage morphology by epifluorescence microscopy. BMDMs were either infected *in vitro* with *L. amazonensis* amastigotes (*iv*LIMs), stimulated with LPS/IFN-γ (M1) or stimulated with IL-4/IL-13 (M2) for three days and compared to untreated/uninfected BMDMs (M0). Analyses were performed using the OPERA Phenix® system. Superpositions of fluorescence images obtained with Hoechst 33342 (nuclear staining, blue) and PhenoVue™ Fluor 488/Concanavalin A (membrane and endoplasmic reticulum staining, green) are shown for one representative experiment out of 10. **B)** Comparative quantitative analysis of M1, M2 and *iv*LIMs secretomes. Violin plot of M0, M1, M2 and *iv*LIMs secretome. Individual scaling was applied to the signal intensity observed for each analyte in the different BMDMs indicated. Subsequently, the signal intensity was subjected to individual scaling for each row. The mean value of every row was computed, and then each intensity in the row was adjusted and transformed into a z-score based on its deviation from the mean (normalized values comprised between −1.5 to +1.5) based on their deviation from the mean. ***, p< 0.001, one-way ANOVA test. **C)** The secretion pattern shared by *iv*LIMs and M1 Mφs (Group 1), *iv*LIMs and M2 Mφs (Group 2), or specific to *iv*LIMs (Group 3) is represented in form of a circular heatmap (n = 3 independent experiments). The fold changes of signal intensity were determined by comparing M1, M2 and *iv*LIMs to M0 Mφs (calibrator). The fold change value was subjected to individual scaling for each row. The mean value of every row was computed, and then each fold change in the row was adjusted and transformed into a z-score based on its deviation from the mean. The brick color represents the z-score of M1, M2 BMDMs and ivLIMs versus M0 BMDMs. **D)** RT-qPCR analysis of a selection of transcripts in short- and long-term *iv*LIMS and in *ev*LIMs. 1) Heatmap representation of the fold expression changes (z-score values) in *iv*LIMs versus M1 and M2 Mφs for 58 selected transcripts quantified by RT-qPCR. Values of the M0 Mφ group were used as calibrators. *iv*LIMs transcripts were classified into three groups according to their similarity of expression to M1 and M2 Mφs: Group 1 (M1-like expression), Group 2 (M2-like expression) and Group 3 (unique expression in *iv*LIMs). Different transcript subgroups were also defined according to their up (a) or down modulation (b). UD, undetected. 2) Analysis of transcript expression levels in day 3, 15 or 30 *iv*LIMs (n = 3 independent experiments) and in *ev*LIMs isolated from footpad lesions of RAG2 KO mice (n = 4 individual mice). Values obtained from mRNA of M0 Mφs were used as calibrator. The grey area corresponds to the range of non-significant expression changes. **E)** Volcano plot showing log2 fold expression changes between *iv*LIMs and uninfected BMDMs. RNA-seq analysis was performed on day 3 post-infection and expression changes between *iv*LIMs and uninfected M0 Mφs were visualized by plotting the Log_2_ fold change in RNA abundance observed between *iv*LIMs vs M0 Mφs (X-axis) and the –Log_10_ adjusted p-value across 3 independent biological replicates (Y-axis). The dots shown in grey correspond to non-significant expression changes (p > 0.05 and −1< Log_2_ fold change < +1). Statistically significant increase or decrease in transcript abundance observed in *iv*LIMs are indicated in red and blue respectively. **F)** Ridgeplots of Gene set enrichment analyses (GSEA) for M1- and M2-related pathways modulated in *iv*LIMs. Differentially expressed genes (RNA-seq) in *iv*LIMs relative to M0 Mφs were subjected to GSEA and the number of genes in each enriched pathway (Y-axis) was plotted against the fold change of core enriched genes for the specified GO term (X-axis). Pathways associated with M1 and M2 polarization are shown in F1 and F2 respectively. The fill color corresponds to the strength of positive or negative enrichment according to the normalized enrichment score (NES).

We obtained first evidenced that *iv*LIMs display a mixt M1/M2 phenotype comparing the secretomes of M1, M2 and infected macrophages using a cytokine array that monitors 111 cytokines, chemokines and acute phase proteins (Figure 1B and Supplementary Figure 2). Integrative analyses of these expression changes revealed a unique secretion profile for *iv*LIMs that overall showed negative Z-scores when compared to the profiles of M1 and M2 polarized BMDMs (Figure 1B). As indicated by the heat map (Figure 1C), *iv*LIMs are characterized by three distinct secretion groups. Group 1 corresponds to an M1-like secretion pattern (19.6% of analyzed proteins), including (i) increased secretion of the pro-inflammatory factors such as WISP1 (Murahovschi et al., 2015) and VEGF, known to recruit monocytes and Mφs (Breen, 2007), or ii) reduced secretion of factors known to dampen *Leishmania*-induced pathology (FLT3L, IL-5, IL-22, MMP9 and EGF) (Gimblet et al., 2015; Kremer et al., 2001; Montoya et al., 2018; Murahovschi et al., 2015; Murase et al., 2018; Watanabe et al., 2004). Group 2 corresponds to an M2-like secretion pattern (38.2% of analyzed proteins), including (i) increased secretion of fibroblast growth factor 21 (FGF-21) known to affect glucose uptake and inhibit macrophage-mediated inflammation (Yu et al., 2016), or (ii) decreased secretion of the anti-leishmanial effectors IL-1α, IL-1β, TNF, IL-12p40, CXCL9 and CXCL10 (Liew et al., 1997; Park et al., 2002; Ritter and Korner, 2002; Schleicher et al., 2016). Group 3 finally corresponds to an *iv*LIMs-specific secretion profile (42.2% of analyzed proteins) characterized by reduced secretion of numerous proteins known to restrict infection, such as IL-33 and RBP4 (pro-inflammatory factors, (Liu et al., 2017; Xu et al., 2019)), CHEMERIN (activator of the inflammasome, (Liang et al., 2019)), CCL2 (activator of the respiratory burst, (Rollins et al.)), PENTRAXIN 2 (activator of the classical complement cascade, (Haapasalo and Meri, 2019)), IL-33, CCL2, AHSG and CD40 (iNOS activators, (Bingaman et al., 2000; Brandonisio et al., 2002; Chattopadhyay et al., 2021; Xu et al., 2019)), sRAGE (driver of the Th1 response, (Chen et al., 2008)) or IL-3, I-L7, G-CSF, and CD40 (factors controlling *Leishmania* elimination, (Brandonisio et al., 2002; Gessner et al., 1993; Ho et al., 1992; Kamanaka et al., 1996; McDowell et al., 2002; Murray et al., 1995; Ritter and Moll, 2000)). Conversely, increased expression in this group was observed for proteins that likely promote parasite survival, including (i) GAS6 (inhibitor of pro-inflammatory cytokine expression, (Alciato et al., 2010)), LEPTIN (inhibitor of NO production and promoter of glucose uptake / glycolytic activity (Becerril et al., 2019; Yang et al., 2019) or VEGF and CD93 (promoters of an anti-inflammatory phenotype (Zhang et al., 2019)).

RT-PCR analysis of 58 polarization markers (Jablonski et al., 2015; Orecchioni et al., 2019; Remmerie and Scott, 2018; Wang et al., 2014) confirmed and extended this mixed phenotype at the transcriptional level, with *iv*LIMs showing expression changes characteristic for M1 polarization (e.g. increased abundance for *ccl3*, *cd68*, *chuk* and decreased abundance for *ccr2*, *mmp9*, *irf4*) and for M2 polarization (e.g. increased abundance for *il1r1*, *dhcr24*, *idi1* and decreased abundance for *ccl2*, *nos2, il1*β, *stat1*) (Figure 1D1). RT-qPCR analysis of 9 selected genes revealed that the mixed profile persisted for at least 30 days post-infection in long-term infected *iv*LIMs and was reproduced in infected tissue macrophages recovered form mouse lesions (*ev*LIMs) thus validating the *in vivo* relevance of our results (Figure 1D2, Supplementary Figure 3A). Both *iv*LIMs and *ev*LIMs further shared unique expression changes when compared to M1 or M2, including (i) increased expression of a series of transcripts implicated in metabolic homeostasis, lipid metabolism and cholesterol biosynthesis (e.g. *igfbp3* (Stuard et al., 2022), *ldlr* (Go and Mani, 2012), *insig1* (Ouyang et al., 2020), *mvd* (Jabalquinto et al., 1988)), and (ii) decreased expression of for example the multifunctional inhibitor of metalloproteases TIMP1 (Grunwald et al., 2019) or pro-inflammatory key actors of the TLR/TNFR/NF-κB pathway, such as *tnfr1a*, *tlr4*, *nf*κ*b1* and *rela*. Furthermore, transcriptional Principal Coordinate Analysis (PCoA) placed *iv*LIMs in between M1 and M2 macrophages (Supplementary Figure 3B), indicating that *Leishmania* orchestrates a secretion profile that avoids activation of its host cells by adopting a M2 profile, while at the same time promoting the recruitment of new monocytes / Mφs via M1-related cytokines/chemokines that likely sustain chronic infection.

For in-depth characterization of the *iv*LIMs immunophenotype, we conducted a comparative transcriptome analysis against M0 Mφs using RNA-seq. *iv*LIMs showed a unique expression profile compared to M0 macrophages, with over 2,300 differentially expressed genes meeting our established criteria for differential gene expression (i.e. Log2 fold change > +1 or < −1, adjusted p-value < 0.05) (Figure 1E). Key genes associated with M1 (*Nos2*) and M2 (*Arg1*) polarization (Das et al., 2010; Munder et al., 1998; Popovic et al., 2007) were significantly upregulated in *iv*LIMs, providing an indication of a mixed M1/M2 phenotype. Gene set enrichment analysis reveals that *iv*LIMs are affected in crucial metabolic pathways linked to M1 (i.e. inhibition of the pentose phosphate pathway or induction of glycolysis) and M2 polarization (i.e. inhibition of fatty acid β oxidation or induction of cholesterol biosynthesis) (Figure 1F1 and 2 respectively and Supplementary Figures 4 and 5).

### *iv*LIMs and *ev*LIMs drive aerobic glycolysis through HIF-1α-independent up-regulation of glycolytic enzymes

RNA-seq analysis revealed that *L. amazonensis* infection may cause an important metabolic shift in *iv*LIMs. Projecting observed expression changes in *iv*LIMs on KEGG maps indeed revealed induction of glycolytic gene expression compared to uninfected M0 (Figure 2A1), suggesting a predominant M1-like energy production. This was further confirmed by RT-qPCR and Western blot analyses showing increased expression levels for *Aldoa*, *Eno1*, *Pkm*, and *Ldha* transcripts (Figure 2A2 and B) and proteins (Figure 2C). Increased expression was sustained during long-term infection *in vitro* (day 15 and 30 *iv*LIMs PI) and validated in infected tissue macrophages recovered from mouse lesions (*ev*LIMs) (Figure 2B). These expression changes were associated with enhanced glycolytic activity, as revealed by increased secretion of L-lactate in *iv*LIMs supernatants, which is exclusively produced by the macrophage as no lactate production was observed in parasite cultures (Figure 2D). Conducting gene set variation analysis based on our RNA-seq results confirmed a global increase in glycolytic gene expression in *iv*LIMs, while genes involved in mitochondrial energy production via oxidative phosphorylation (OXPHOS) or the Tricarboxylic Acid (TCA) cycle were either not modulated (OXPHOS) or down-regulated (TCA) (Figure 2E, Supplementary Figures 6 and 7).

**FIGURE 2.**
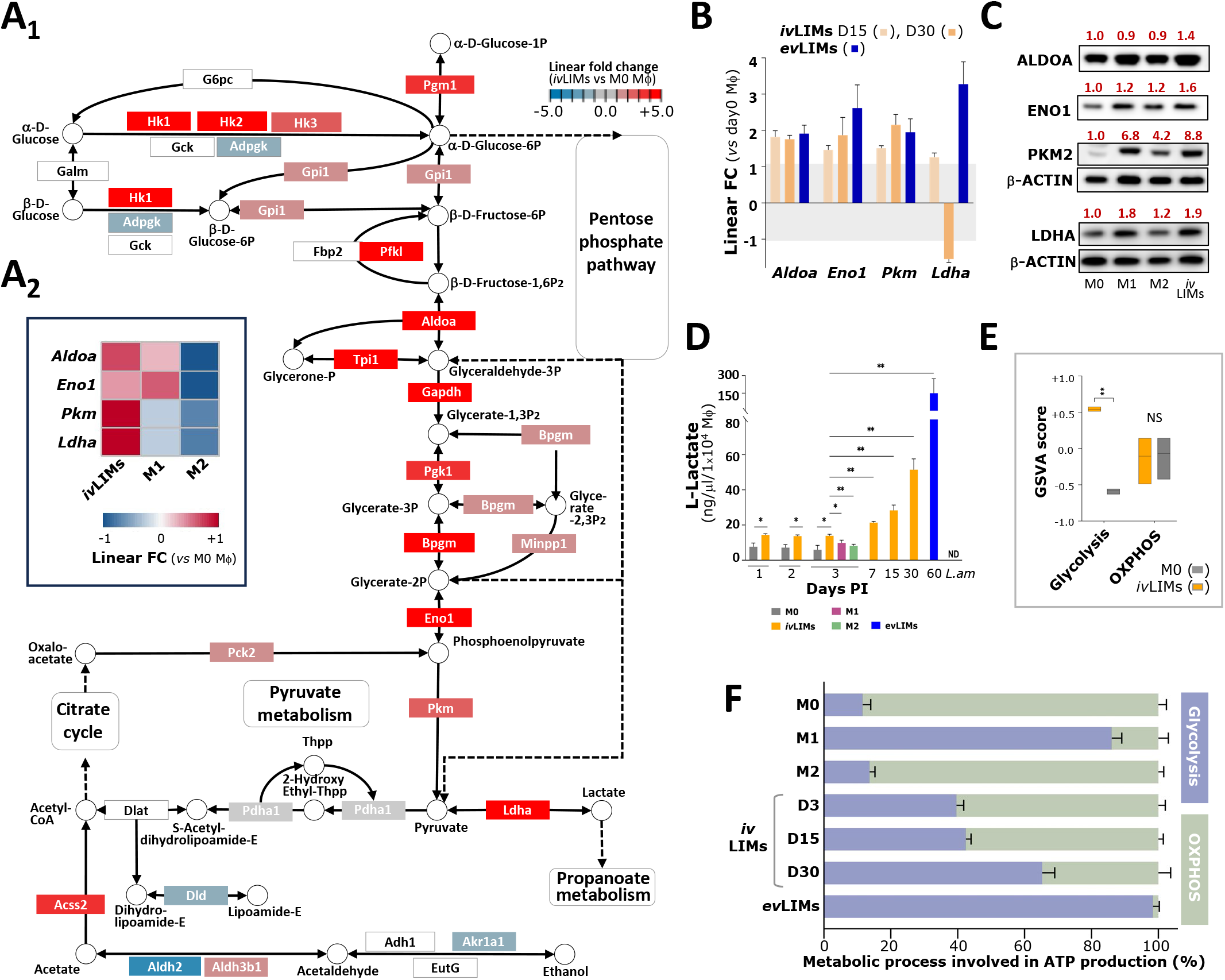
Analysis of glycolytic enzymes showing differential transcript abundance in *iv*LIMs. **A1)** Projection of expression changes in *iv*LIMs compared to M0 BMDMs onto the KEGG map of the glycolytic pathway. Increased and decreased mRNA abundances in *iv*LIMs compared to M0 are respectively indicated in red and blue intensities according to the shown legend. **A2)** Transcript abundance of the indicated glycolysis genes was determined in *iv*LIMs, M1, M2 and M0 Mφs by RT-qPCR analysis and expression changes are represented as a heatmap using the values of M0 Mφs as calibrator. Increased and decreased mRNA abundances compared to M0 are respectively indicated in red and blue intensities according to the shown legend. **B)** Transcript abundance of the indicated glycolysis genes was determined in *iv*LIMs at day 15 and day 30 post-infection and in tissue-isolated, infected macrophages (*ev*LIMs) by RT-qPCR analysis and linear fold changes (FC) were determined in 3 independent biological replicates. The grey area corresponds to non-significant FC values (p > 0.05). **C)** Western blot analysis of M0, M1, M2 BMDMs and *iv*LIMs extracts obtained at day 3 PI for aldolase A (ALDOA), enolase 1 (ENO1), type M2 pyruvate kinase (PKM2), lactate dehydrogenase A (LDHA) and β-ACTIN. Signals were quantified using the ImageQuant imaging system utilizing β-ACTIN as calibrator. Red numbers indicate the ratio of band intensity relative to M0 expression after normalization against β-ACTIN. **D)** Determination of L-lactate levels in supernatants from M0, M1, M2, *iv*LIMs and *ev*LIMs after passing 10 kDa cut-off filter by centrifugation to remove lactate dehydrogenase (LDH). Subsequently, the lactate concentration in each supernatant was assessed using an L-lactate assay kit and normalized based on cell number (n = 3 independent experiments). *, p < 0.05; **, p < 0.01, one-way ANOVA test. **E)** Gene set variation analysis (GSVA) for the OXPHOS and glycolysis pathways based on the normalized read counts of the RNA-seq data. The global regulation of pathways in each sample were computed based on GSVA statistic methods. **, p < 0.001, t-test. **F)** ATP production rate dependent on OXPHOS (violet bar) or glycolysis (grey bar) was monitored applying the Agilent Seahorse XF Real-Time ATP Rate Assay Kit on M0, M1 and M2 Mφs, *iv*LIMs at day (D) 3, 15 and 30 PI, and tissue-isolated, infected macrophages (*ev*LIMs). N = 3 biological replicates.

We next validated this metabolic shift in energy production in live cells determining glycolytic and OXPHOS ATP production rates *in vitro* in short- and long-term cultures of *iv*LIMs and *ex vivo* in *ev*LIMs through real-time metabolic analysis. We capitalized on the absence of the typical complex I of the respiratory chain in *L. amazonensis* (Duarte et al., 2021; Nebohacova et al., 2009) to dissociate the host and parasite contributions to the measured Oxygen Consumption Rate (OCR) comparing the effects of the inhibitors rotenone (complex I inhibitor, targets only host OXPHOS) and antimycin A (complex III inhibitor, targets both host and parasite OXPHOS) alone or in combination. Indeed, free amastigotes produced a significant OCR signal, which contributed more than 40% to the basal signal observed in *iv*LIMs (Supplementary Figure 8). Assessing the net glycolytic and OXPHOS activities in M0, M1 and M2 polarized macrophages, in *iv*LIMs from short-, mid- and long-term infections as well as in *ev*LIMs revealed a progressive increase in glycolytic-versus OXPHOS-dependent ATP production that attained 98% in lesion-derived *ev*LIMs and thus levels even exceeding M1 polarized BMDMs (86.1%) (Figure 2F).

Collectively, these results reveal that infected LIMs favor aerobic glycolysis (also known as the Warburg effect) for ATP production through the coordinated upregulation of transcripts encoding key glycolytic enzymes from the early stages of infection to long-term chronic infections both *in vitro* and *in vivo*.

### The increase in aerobic glycolysis in *iv*LIMs is independent of HIF-1α up-regulation

The increased expression of glycolytic enzymes during hypoxic or normoxic conditions is primarily regulated by induction of the transcription factor (TF) HIF-1α (Kierans and Taylor, 2021; Vaupel and Multhoff, 2021), but also other TFs such as c-MYC, K-RAS, STAT3, TRP53, ZEB1, OCT1, or ERG (Jiang et al., 2022; Li et al., 2021; Yeung et al., 2008; Zhang et al., 2020). Unlike expected from the Warburg effect we observed in *iv*LIMs, the elevated expression of glycolytic genes in infected macrophages did not correlate with increased expression of these TFs, which predominantly showed reduced expression (Figure 3A). In particular, unlike M1 macrophages, no significant expression changes were detected at mRNA and protein levels for HIF-1α in *iv*LIMs and lesion-derived *ev*LIMs (Figure 3B and C, left panels). Overall, these findings suggest that neither HIF-1α nor other TFs contribute to the upregulation of glycolytic enzymes we observed in *iv*LIMs. We therefore investigated alternative regulatory mechanisms known to control the expression of glycolytic enzymes.

**FIGURE 3.**
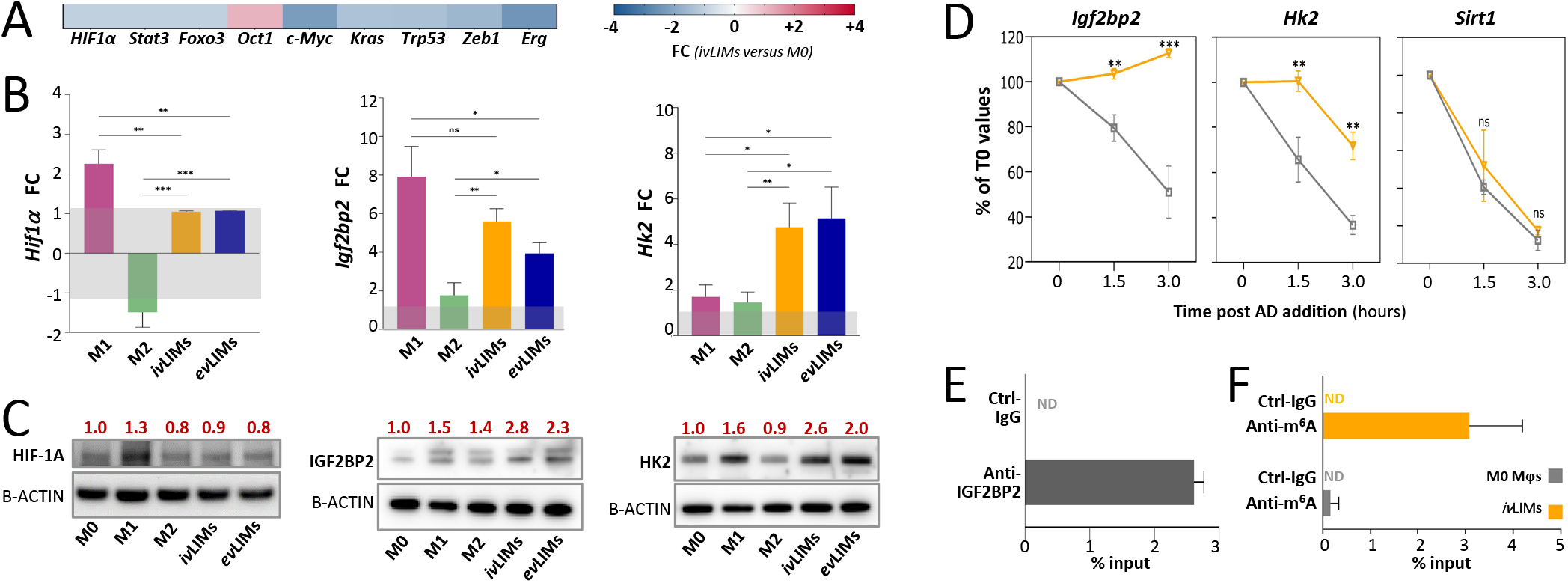
Post-transcriptional regulation of *HK2* mRNA in *iv*LIMs. **A)** Fold changes in mRNA abundance (RNA-seq) observed in *iv*LIMS compared to M0 Mφs for TFs known to regulate glycolysis genes. **B)** RT-qPCR analysis of total RNA extracts from M0, M1, M2 Mφs, *iv*LIMs and *ev*LIMs for *HIF-1*α*, Igf2bp2* and *Hk2* transcripts. M0 was used as a calibrator to calculate fold changes. The grey area corresponds to non-significant FC values (p > 0.05). *: p < 0.05; **, p < 0.01; one-way ANOVA test; n = 3 independent experiments. **C)** Western blot analysis of HIF-1α, IGF2BP2 and HK2 using total cellular extracts of M0, M1, M2 Mφs, *iv*LIMs and *ev*LIMs. Signals were quantified using the ImageQuant imaging system utilizing β-ACTIN as calibrator. Red numbers indicate the ratio of band intensity relative to M0 expression after normalization against β-ACTIN. **D)** Evaluation of mRNA stability in *iv*LIMs and M0 Mφs of *Igf2bp2, Hk2 and Sirt1* transcripts following Actinomycin D-treatment. RNA was isolated at 0, 1.5 and 3 hours after treatment and subjected to RT-qPCR analysis. Fold change expression ratios were determined by comparison to untreated Mφs at t = 0h. **, p<0.01; ***, p<0.001; t-test; n = 3 independent biological replicates; ns, no significant differences. **E)** Quantification of IGF2BP2-bound *Hk2* transcript by RNA immunoprecipitation (RIP) in protein lysates of M0 Mφs using anti-IGF2BP2 and negative control IgG antibodies (Ctrl-IgG). After elution of bound mRNA, *Hk2* transcripts were quantified by RT-qPCR. Results represent the mean and SEM of the percentage of input. n = 3 independent biological replicates; ND, transcripts not detected. **E)** Quantification of m^6^A modification levels in M0 Mφs (grey) and *iv*LIMs (orange) of *Hk2* transcripts by Methylated RNA immunoprecipitation (MeRIP) using m^6^A-RNA-specific or control IgG antibodies samples. Mean values and SEM of percentage of input is shown. n = 3 independent biological replicates; ND, transcripts not detected.

Aerobic glycolysis (Warburg effect) is a hallmark of cancer cells that often correlates with HIF-1α induction, but also involves post-transcriptional regulation of glycolytic transcripts via RNA-binding proteins (RBPs) (Wegener and Dietz, 2022). We therefore mined or RNA-seq data for expression changes of RBPs that are predicted to bind to glycolytic transcripts using the POSTAR3 database. We identified 11 glycolytic transcripts with increased abundance in *iv*LIMs that are targets of 44 RBPs (Supplementary Figure 9). The RBP insulin-like growth factor 2 mRNA-binding protein 2 (IGF2BP2) attracted our attention for several reasons: (i) it showed increased expression in both *iv*LIMs and *ev*LIMs at mRNA and protein levels and thus correlates with expression changes of most glycolytic enzymes (Figure 3B and C, middle panels), (ii) it is predicted to bind the mRNA of 9 out of the 11 glycolytic transcripts induced in *iv*LIMs (Supplementary Figure 9), and most importantly (iii) it serves as a m^6^A-RNA reader protein enhancing mRNA stability of hexokinase 2 (*HK2*), which is a key mediator of aerobic glycolysis (Liu et al., 2021; Xu et al., 2022).

To further corroborate post-transcriptional induction of glycolysis in *iv*LIMs, we directly investigated differential mRNA turn-over in M0 BMDMs and *iv*LIMs. Cells were treated with the transcription inhibitor actinomycin D (ActD) and mRNA was monitored by RT-PCR at 1.5 and 3h post-treatment for IGF2BP2, its putative target *Hk2,* and *Sirt1* as a negative control. While no significant changes were observed for *Sirt1,* the half-life of *Hk2* mRNA was significantly increased in *iv*LIMs compared to M0 BMDMs, confirming post-transcriptional regulation of this glycolytic enzyme (Figure 3D). Surprisingly, *Igf2bp2* itself is post-translationally regulated in *iv*LIMs as judged by the absence of mRNA degradation during the 3h experimental window, suggesting a positive, auto-regulatory feedback loop. Combining RNA immunoprecipitation (RIP) with RT-qPCR, we next revealed a direct interaction of *Igf2bp2* with its *Hk2* mRNA in M0 BMDMs, leading to a possible stabilization of this target mRNA and its increased abundance we observed in *iv*LIMs (Figure 3E). This possibility was further supported by methylated mRNA immunoprecipitation (MeRIP) that showed increased m^6^A-RNA methylation levels of *Hk2* mRNA in *iv*LIMs compared to M0 BMDMs, thus increasing the binding affinity for the m^6^A reader IGF2BP2 (Figure 3F).

In conclusion, our data suggests a HIF-1α-independent, alternative pathway of metabolic reprogramming in *iv*LIMs that likely relies on epi-transcriptomic control of glycolytic gene expression via a regulatory interaction between IGF2BP2 and m^6^A methylated *Hk2* mRNA resulting in increased transcript stability and abundance.

### The m^6^A RNA reader IGF2BP2 is required for induction of aerobic glycolysis in *iv*LIMs

To functionally validate the role of IGF2BP2 in the epi-transcriptomic control of *Hk2* mRNA abundance and aerobic glycolysis in *iv*LIMs, we assessed the impact of siRNA-mediated silencing of *Igf2bp2* on glycolytic gene expression. After establishing optimal conditions for siRNA transfection (Supplementary Figure 10), we successfully reduced IGF2BP2 expression as judged by 80% and 50% reduction of mRNA and protein abundances respectively by two of the three transfected siRNAs (Figure 4A and B). A significant decrease in *Hk2* transcript abundance was observed in these cells, further sustaining the IGF2BP2/*Hk2* regulatory relationship (Figure 4A). *Igf2bp2* knockdown had no significant effect on the oxygen consumption rate (OCR, Figure 4C1) in *iv*LIMs, suggesting that the level of OXPHOS-dependent ATP production is not regulated by this factor. In contrast, the knockdown of *Igf2bp2* resulted in a notable decrease in the glycolysis (35.5% reduction for si-*Igf2bp2*-1) and glycolytic capacity (34.6% for si-*Igf2bp2*-1) in response to glucose and oligomycin addition (Figure 4C2 and 3). This observation confirms the direct involvement of this m^6^A reader protein in enhancing glycolytic activity in LIMs, possibly through the stabilization of *Hk2* mRNA.

**FIGURE 4.**
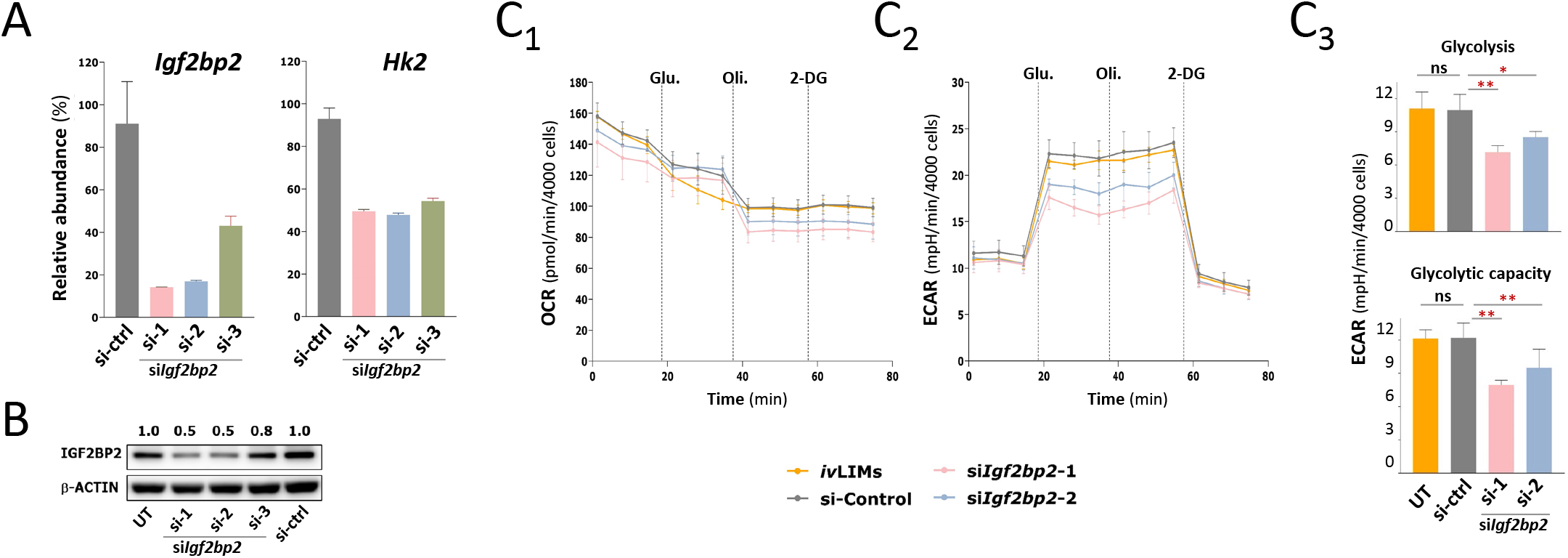
Impact of *Igf2bp2* knock down in *iv*LIMs. **A)** RT-PCR analysis. Three distinct small interfering (si) RNAs (si-1, −2, −3) targeting *Igf2bp2*, along with a universal negative control siRNA (si-ctrl), were assessed in M0 Mφs on the first day post-differentiation at a concentration of 50 nM. The transcripts of *Igf2bp2* and of the IGF2BP2 target *Hk2* in untransfected and transfected M0 Mφs 24 hours post-transfection were quantified by RT-qPCR (n = 3 independent experiments). **B)** Western blot analysis. Cell lysates from untransfected and transfected M0 Mφs using 50 nM of the indicated siRNAs (si-1, −2, −3) and universal negative control siRNA (si-ctrl) were subjected to Western blot analysis to assess the expression of IGF2BP2 and β-ACTIN (normalization control) at 48 hours post-transfection. The knock down efficiency was assessed by quantifying the IGF2BP2 signals using the ImageQuant imaging system. Numbers above the lanes indicate the ratio of band intensity relative to untransfected M0 Mφs after normalization against the β-ACTIN signal. Due to their higher efficiency, only si-1 and si-2 were be retained for further experiments. **C)** Seahorse analysis of the effect of *Igf2bp2* knockdown on *iv*LIMs glycolysis. The Seahorse glycolysis stress test was performed on *iv*LIMs one day following siRNA transfection (10 nM). Glucose, oligomycin, and 2-Deoxy-D-glucose (2-DG) were sequentially added as indicated. The Oxygen Consumption Rates (OCR) and extracellular ECAR acidification rate (ECAR) were measured for *iv*LIMs with or without siRNA treatment. Results were normalized by cell number. Glycolysis and glycolytic capacity of *iv*LIMs with or without siRNA treatment were calculated. Results were normalized by cell number. ns, no significant difference; *, p < 0.01; **, p < 0.001, one-way ANOVA test. N = 3 independent experiments.

## Discussion

We used a multiparametric, immuno-metabolic profiling approach to characterize in-depth the phenotype of *Leishmania*-infected, primary macrophages *in vitro* and lesional macrophages *ex vivo* (respectively termed *iv*LIMs and *ev*LIMs). We uncovered a complex, atypical polarization state in these LIMs that express both M1 and M2 polarization markers, while favoring an M1-like profile of energy production relying on aerobic glycolysis. We showed that this form of energy production was independent of HIF-1α-mediated regulation in LIMs but instead controlled at epi-transcriptomic level by the m6A reader enzyme IGF2BP2 through direct binding and stabilization of of m6A-modified *Hk2* transcripts. Our results challenge the prevailing notion that intracellular *Leishmania* simply triggers an M2-like phenotype while avoiding the expression of M1 marker genes (Das et al., 2021; Kumar et al., 2018; Lecoeur et al., 2020; Roy and Mandal, 2016) and open important new questions on (i) the relevance of the LIMs mixed polarization profile for clinical infection, (ii) the roles of aerobic glycolysis in promoting intracellular parasite survival, and (iii) the mechanisms of HIF-1α-independent regulation of the LIMs energy metabolism.

Macrophages represent a set of highly divergent cells that adopt different phenotypic characteristics according to developmental stage, tissue environment, immune challenge or disease state (Davis et al., 2013). For example, tumor-associated macrophages (TAMs) promote a microenvironment beneficial for cancer metastasis, which has been associated with the expression of a mixed M1/M2 polarization profile (Hambardzumyan et al., 2016). Reminiscent to TAMs, our study demonstrates that the protozoan parasite *Leishmania* establishes a complex polarization profile in infected host macrophages that defines yet another type of disease-associated macrophages we termed LIMs (for *Leishmania*-infected macrophages). As observed for TAMs that show important variations according to the type of cancer they are associated with (Garrido-Martin et al., 2020; Pe et al., 2022), LIMs too likely show phenotypic variability according to infecting parasite species, host immune status, or the type of infected tissue. In our experimental system of cutaneous leishmaniasis using *L. amazonensis* lesion-derived amastigotes, LIMs adopt a M1/M2 mixed polarization profile and express LIMs-specific markers, which together reveal a dual subversion strategy that triggers an anti-inflammatory response in *cis* inside infected host cells thus promoting parasite survival, while at the same time driving a TH1 response in *trans* promoting cell recruitment that can amplify the infection (Figure 5). This dual macrophage subversion strategy echoes the dual Th1/Th2 response observed in experimental visceral (VL) and cutaneous leishmaniasis (CL) caused by *L. donovani* and *L. panamnensis*, respectively (Melby et al., 1994; Osorio et al., 2003), in symptomatic dogs infected with *L. infantum* (de Almeida Leal et al., 2014), or human VL patients (Nylen and Sacks, 2007).

**FIGURE 5.**
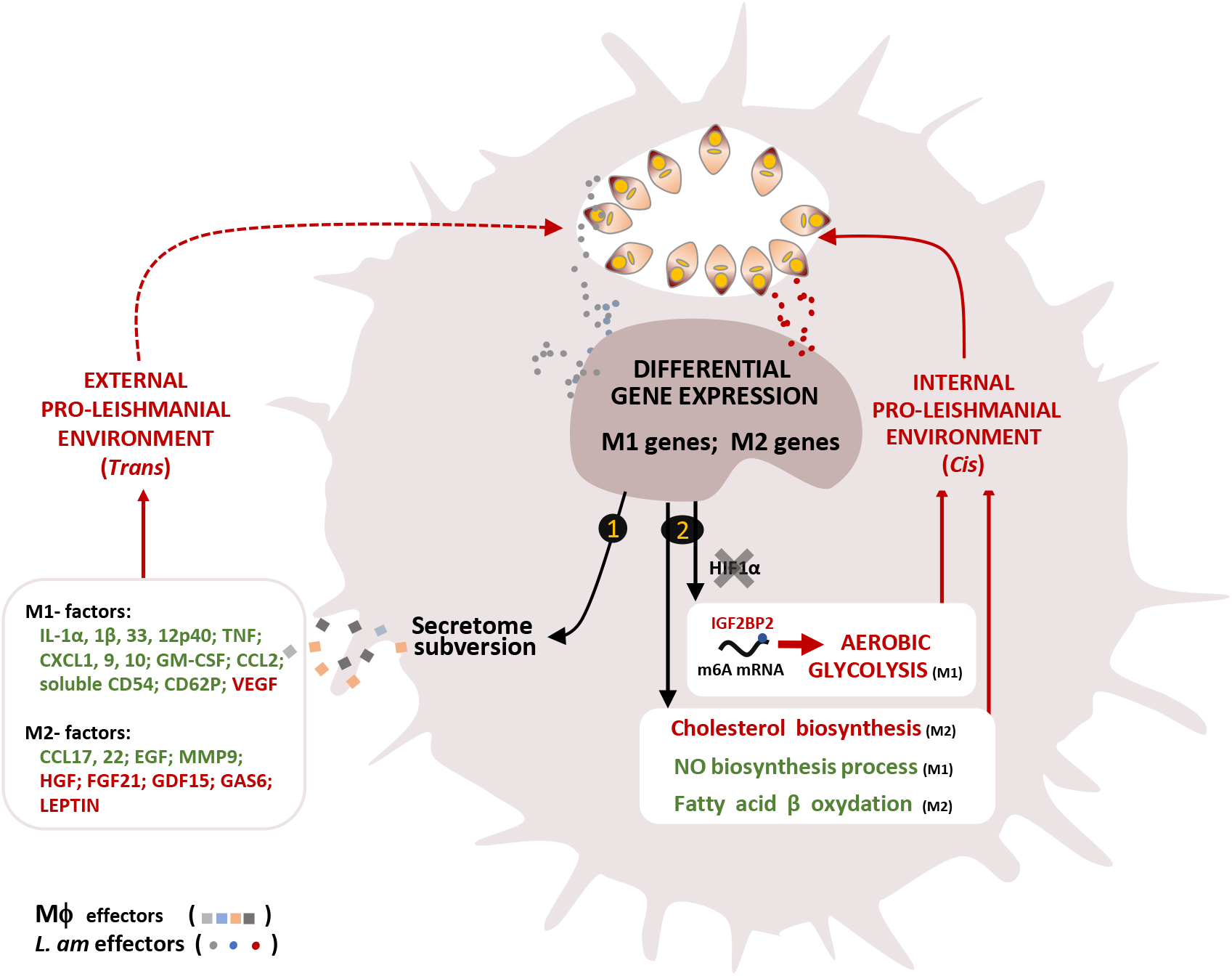
*L. amazonensis* induces a unique immuno-metabolic phenotype of its host macrophage. Intracellular *L. amazonensis* establish a characteristic gene expression pattern of their host macrophage, probably through the secretion of effectors / metabolites that remain to be defined. These expression changes indicate remodelling of both immune (path 1) and metabolic (path 2) features of the infected macrophage to generate external and internal environments permissive for parasite and host cell survival. Path 1: Induction of a permissive immunological environment. Parasites remodel the macrophage secretome (cytokines, chemokines and acute phase proteins) towards a unique pattern of M1-like, M2-like and LIMs-specific secreted factors. Factors that show decreased or increased secretion in LIMs are displayed in green and red, respectively. This unique LIMs’ secretome likely abrogates Mφ activation, favors intracellular survival / multiplication and prevents the development of an anti-leishmanial Th1 response *in vivo*. Path 2: Induction of a permissive metabolic phenotype. LIMs display a unique metabolism characterized by both M1-features (e.g. inhibition of NO synthesis, increased glycolysis) and M2-features (e.g. inhibition of fatty acid β oxidation, increase of glutamine metabolism). LIMs shift ATP production from mitochondrial OXPHOS to aerobic glycolysis, which occurs in a HIF-1α independent, IGF2BP2-dependent manner. These metabolic changes may again favor both parasite and macrophage survival.

Macrophage polarization is tightly connected to metabolic reprogramming, with M1 macrophages relying on aerobic glycolysis for energy production, while M2 favor mitochondrial oxidative phosphorylation for ATP production (De Santa et al., 2019). Surprisingly, despite their mixed M1/M2 polarization profile, there was a clear bias of LIMs towards aerobic glycolysis, with infected, lesional macrophages relying nearly exclusively on aerobic glycolysis for energy production. Our results establish a first assessment of the LIMs energy metabolism during chronic infection *in vitro* and *in vivo* and call into question previous observations that linked *in vitro* short-term infection to induction of mitochondrial energy production linked to increased macrophage mitochondrial biogenesis (Acevedo Ospina et al., 2022; Moreira et al., 2015). It remains to be established whether this discrepancy relies on the different infection systems used or low infection levels monitored in these reports.

Our findings raise the question to what extent this metabolic reprogramming benefits the parasite. Aerobic glycolysis - also known as the "Warburg effect" - was initially defined as a metabolic process displayed by tumor cells under normoxic conditions, where glucose is oxidized into pyruvate and then converted to lactate (Warburg et al., 1927) despite the presence of oxygen. While this metabolic profile contrasts with the more energy efficient OXPHOS pathway observed in resting, non-cancer cells (Schurr and Passarella, 2022), it mobilizes energy much faster thus supporting cancer cell proliferation. Give that macrophages are terminally differentiated and that LIMs do not proliferate, this Warburg effect may rather establish conditions that could promote parasite survival, such as evasion of anti-microbial ROS production generated by OXPHOS (Liemburg-Apers et al., 2015), or sustained long-term survival of infected host cells, a phenomenon we previously documented in *iv*LIMs and *ev*LIMs (Lecoeur et al., 2022). Indeed, the production of lactate during aerobic glycolysis does not simply represent a metabolic waste product but has been identified as a major biological regulator that can (i) counteract pro-inflammatory cytokine production (Yang et al., 2020), (ii) participate in macrophage recruitment (Chen et al., 2020), (iii) enhance transcription of immunosuppressive IL-10 (Heim et al., 2020), and (iv) confer protection to different forms of cell death (Huang et al., 2013; Ma et al., 2022; Pradelli et al., 2010; Sanman et al., 2016; Vaughn and Deshmukh, 2008).

Finally, one of the key findings of our report is represented by HIF-1α-independent induction of aerobic glycolysis. This form of energy production is usually induced by HIF-1α that can show increased expression both under hypoxic or normoxic conditions and acts as a major transcriptional activator glycolytic enzymes. Surprisingly, even though there is ample evidence for an important role of HIF-1α induction during infection with various *Leishmania* species (Alonso et al., 2019; Arrais-Silva et al., 2005; Hammami et al., 2017; Hammami et al., 2015; Mesquita et al., 2020; Rocha et al., 2018; Weinkopff et al., 2019), expression and protein abundance of this TF were neither altered in *iv*LIMs nor *ev*LIMs, suggesting alternative forms of regulation. While we cannot rule out possible post-translational modifications that may alter HIF1-α TF activity (Albanese et al., 2020), we identified the m^6^A reader protein IGF2BP2 as a possible key regulator underlying HIF-1α-independent LIMs metabolic reprogramming. This mRNA-binding protein is known to control glycolytic metabolisms by stabilizing various transcripts of the glycolytic pathway, including m^6^A modified *Ldha*, *AldoA* or *Hk2* transcripts (Liu et al., 2021; Wu et al., 2023; Zhou et al., 2024). Here we report that a similar scenario applies for LIMs as judged by (i) increased IGF2BP2 expression, (ii) increased abundance of 9 established IGF2BP2 target mRNAs involved in glycolysis; and (iii) increased m^6^A modification of the key target *Hk2 mRNA.* While these results are only correlative in nature, we firmly established a role of IGF2BP2 in LIMs aerobic glycolysis by demonstrating enhanced IGF2BP2/*Hk2* mRNA interaction in *iv*LIMs and significant reduction of glycolytic activity and *Hk2* mRNA abundance following *Igf2bp2* knock down.

In conclusion, our study uncovers a surprisingly complex and fine-tuned “à la carte” phenotypic shift in LIMs that rules out a simple activation of just the M2 polarization program. The unique immuno-metabolic homeostasis of LIMs establishes both intracellular and extracellular conditions for parasite survival and chronic infection, which resonates with the regulatory Mφs linked to human diffuse cutaneous leishmaniasis (Christensen et al., 2019), or the dual Th1/Th2 response observed during experimental/clinical leishmaniasis (Melby et al., 1994; Osorio et al., 2003). Our results reveal for the first time a direct and sustained effect of these parasites on the host cell energy-producing pathways, and define IGF2BP2-dependent, epi-transcriptomic regulation as a novel strategy for parasite-mediated host cell subversion.

## SUPPLEMENTARY MATERIAL AND METHODS

### Isolation, culture, and seeding of bone marrow-derived macrophages

Bone marrow cells were recovered from tibias and femurs of C57BL/6 mice and cultured in bacteriological Petri dishes in complete DMEM medium (Gibco Life Technologies, 2140821) containing 10% FCS, 50 units/µg/ml of penicillin/streptomycin and 50 µM β-mercaptoethanol and supplemented with 50 ng/ml of mouse recombinant colony-stimulating factor 1 (rm-CSF-1, ImmunoTools GmbH, Germany, 682087) (Lecoeur et al., 2020). After 6 days at 37°C in a 7.5% CO_2_ atmosphere, adherent Mφs were detached with 25 mM EDTA in PBS, and seeded in a complete medium into different plastic cell culture supports, including (i) tissue culture-treated Petri dishes (Falcon® 100 mm TC-treated Cell Culture Dish, OPTILUX 353003) for RNA-seq, (ii) 384-well plates (culture-treated, flat-optically clear bottom, Cell carrier plate, PerkinElmer) for morphological analyses on images acquired with the OPERA Phenix automated spinning disk confocal imager, (iii) tissue culture-treated 6-well plates (Corning Life Science, 353046) for RT-qPCR and Western blot analyses, and iv) in Seahorse XFe96 tissue culture microplates (Agilent, 102601-100) for metabolic flux analyses.

### Quantification of nitrites

M0, M1-like, M2-like or *iv*LIMs were cultured in tissue culture-treated 6-well plates (Corning Life Science, 353046) at 34°C. After 3 days of incubation, supernatants were collected and filtered using a 0.2 μm disposable filter (Gamme ClearLine, 146564). Next, 20 μL of Griess reagent colorimetric Griess test kit (Thermo Fisher Scientific, G7921) (equal volumes of N- (1-naphthyl) ethylenediamine and sulfanilic acid mixture), 150 μL of supernatant and 130 μL of deionized water were mixed and incubated for 30 minutes in a 96-well plate (Corning, 353072) at room temperature (RT). Nitrite quantitation was based on the measurement of the absorbance at 548 nm of each sample and calculated using a standard curve.

### Western Blotting (WB)

9 μg of total protein extracts was loaded onto each lane on NuPAGETM Bis-Tris gels (ThermoFisher Scitentific, NP0326BOX), and resolved in NuPAGETM MOPS (ThermoFisher Scitentific, NP0001) or NuPAGE^TM^ MES SDS running buffer (ThermoFisher Scitentific, NP0002). Proteins were electro-transferred onto a 0.45 μm PVDF membrane (ThermoFisher Scientific, 88518) at 4°C. Membranes were blocked for 1 hour at RT using the ECL Prime Blocking Reagent (Cytiva, RPN418). Protein expression levels were detected by incubation with primary antibodies specific for ALDOA (11217-1-AP), ENO1 (11204-1-AP), PKM2 (15822-1-AP), LDHA (19987-1-AP), IGF2BP2 (11601-1-AP), HK2 (22029-1-AP) B-ACTIN (20536-1-1AP) and HIF-1α (20960-1-AP) all provided by Proteintech, followed by incubation with secondary antibodies conjugated with horseradish peroxidase (ThermoFisher Scientific, 31462). β-ACTIN expression was assessed as a normalization control for protein loading. Protein bands were visualized with the ECL Prime reagent (Cytiva, RPN2232), followed by imaging with the ImageQuant^TM^ 800 system (Cytiva), and quantification with the ImageQuantTL software (v10.1-401). HIF-1α analysis revealed several bands as previously published (Bera et al., 2019; Liu et al., 2020a).

### Proteome Profiler Mouse XL Cytokine Array

Supernatants of Mφ cultures from 3 independent biological replicates were collected, pooled, and added to the membranes provided by the Proteome Profiler Mouse XL Cytokine Array (ARY028, Biotech, R&D systems) according to the manufacturer’s instructions. Luminescence detection and quantification were performed by the ImageQuantTL software. Protein levels were expressed as Z-scores based on the relative expression of the M0 group (Liu et al., 2020b). Scripts for Heatmap and Venn diagrams of Z-scores were based on native graphical functions and run using R version 4.1.2.

### Sample preparation or morphologic analysis of BMDM cultures

Fixed BMDMs were subjected to nuclear staining with Hoechst 33342, and to cellular membranes/endoplasmic reticulum staining with PhenoVue™ Fluor 488-Concanalin A (Perkin Elmer). Neutral lipid droplets were stained using BODIPY™ 493/503 and/or Diphenylhexatriene (DPH; ThermoFisher Scientific). Acquisition of bright-field and fluorescence images was carried out using the confocal and non-confocal modes with a 40x water objective. Subsequent analysis of the acquired images was performed using the Columbus™ Image Data Storage and Analysis System (PerkinElmer).

### Determination of the bioenergetic profile of macrophages

Samples were analyzed by the Seahorse XFe96 analyzer according to the manufacturer conditions. ATP production via glycolysis and mitochondrial oxidative phosphorylation was determined by respectively measuring Extracellular Acidification Rate (ECAR) and Oxygen Consumption Rate (OCAR) at 37°C in Seahorse XF DMEM medium (pH=7.4, with 10 mM glucose, 2□mM L-glutamine, and 1□mM sodium pyruvate) (Agilent, 103680-100) as defined in the Agilent user guide. Oligomycin (1.5 μM), an inhibitor of ATP synthase in the complex V of the mitochondrial electron transport chain (ETC), was injected into each well after 18 minutes of measurement, then 0.5 μM of the inhibitors Rotenone (ROT, inhibitor of complex I of the ETC) and Antimycin A (AA, inhibitor of complex III of the ETC) alone or as a mixture were injected after 36 minutes of measurement. ATP-related parameters were also measured, including the rate of ATP production associated with i) the conversion of glucose to lactate in the glycolytic pathway (GlycoATP production rate), and ii) the oxidative phosphorylation in the mitochondria (mitoATP production rate) using oxygen consumption, as a read out. The total ATP production rate was calculated according to the manufacturer’s instructions and normalized by cell number per well (Counting of Hoechst 33342-stained nuclei).

For the analysis of the mitochondrial function of macrophages and amastigotes (Agilent Seahorse XF Cell Mito Stress Test Kit), BMDMs, amastigotes and *ev*LIMs were plated and cultured as described above and a series of injections was performed sequentially into each well of the system. At the 18 minute, 1.5 µM of Oligomycin was introduced, followed by the sequential injection of the uncoupler Carbonyl cyanide-4 phenylhydrazone (FCCP, 1 μM) at 36 minutes. Finally, 0.5 µM of either individual inhibitors (ROT or AA) or a combination of (Rot-AA) was injected after 54 minutes. OCR and ECAR normalization per well were performed after completion of the analysis by counting Hoechst 33342-stained macrophages on pictures (taken at 4X using the EVOS microscope) using the ImageJ software (version 1.54g).

### RT-qPCR analysis

After sample lysis, genomic DNA was removed using a DNA removal column provided by the kit. Binding solution was added to the eluate, mixed thoroughly, deposited on an RNA retention column and total RNA (>= 200 nucleotides) was eluted. RNA quantity and purity was assessed using the NanoDrop (ThermoFisher Scientific, NanoDrop^TM^ 1000 Spectrophotometer) and stored at −20°C. RT-qPCR was carried out in 384-well PCR plates (Framestar 480/384, 4titude, Dominique Dutscher) using the iTaq Universal SYBR® Green Supermix (Bio-Rad) and 0.5 μM primers with a LightCycler® 480 system (Roche Diagnostics, Meylan, France). Primer information for all targets is detailed in the Supplementary Table S3. Crossing Point values were determined using the second derivative maximum method (LightCycler® 480 Basic Software). The relative expression software tool (REST©-MCS) (Pfaffl et al., 2002) was used to calculate fold change (FC) values. Normalization was performed with the geometric mean of the two best reference genes *Ppih* and *Mau2* as determined by the Normfinder programs (data not shown). For statistical analysis of gene expression levels, Cp values were first transformed into relative quantities (RQ) and normalized (Lecoeur et al., 2020). Nonparametric Kruskal-Wallis tests were performed on Log-transformed Normalized Relative Quantity values. Heatmap were generated (R, version 4.1.2) using homemade scripts based on native graphical functions. The fold change value was subjected to individual scaling for each row. The mean value of every row was computed, and then each fold change in the row was adjusted and transformed into a z-score based on its deviation from the mean. Results were expressed as z-scores calculated by relative expression compared to the M0 group.

### RNA-seq analysis

RNA samples were treated with DNAse and then processed for library preparation using the TruSseq Stranded mRNA sample preparation kit (Illumina France, Evry), according to the manufacturer’s instructions. To selectively isolate the polyadenylated RNA fraction and remove ribosomal RNA, an initial poly(A) RNA isolation step was performed on 1 µg of total RNA, as outlined in the Illumina protocol. Subsequently, the enriched fraction underwent fragmentation using divalent ions at high temperatures. The fragmented RNA samples were then subjected to random priming for reverse transcription, followed by second-strand synthesis without end repair step, to generate double-stranded cDNA fragments. An adenine residue was added to the 3’ end of the cDNA fragments, and specific Illumina adapters were ligated. After PCR amplification of the ligation products, the oriented libraries were assessed using Bioanalyzer DNA1000 Chips (Agilent, # 5067-1504) and quantified using spectrofluorometry (Quant-iT™ High-Sensitivity DNA Assay Kit, #Q33120, Invitrogen). Sequencing was performed on the Illumina Hiseq2500 platform, generating single-end 65 bp reads with strand specificity.

The obtained reads were cleaned using cutadapt (version 1.11) (Kechin et al., 2017), and only sequences with a minimum length of 25 nt were employed for further analysis. Reference genome alignment (GRCm38 from Ensembl database 94) was performed using STAR (version 2.5.0a) (Dobin et al., 2013) with default parameters. FeatureCounts (version 1.4.6-p3) (Liao et al., 2014) from the Subreads package (Tugal et al., 2013) was used for Gene counting, with the parameters set to count genes and consider both strands. The transcriptomic data generated in this study have been deposited in the NCBI’s Gene Expression Omnibus repository under the subseries GSE205856 and are publicly available for further analysis and mining.

### RNA immunoprecipitation (RIP)

Adherent M0 or *iv*LIMs were washed twice with 10 mL of ice-cold PBS, collected by scraping and pelleted by centrifugation (450 g, 5 minutes at °C). The supernatant was discarded and 5×10^7^ cells were homogenized and lysed with 250 μL of Polysome Lysis Buffer (PLB) containing 100 mM KCl (Sigma-Aldrich, P391), 5 mM MgCl_2_ (Sigma-Aldrich, M9272), 10 mM HEPES-NaOH pH 7.0 (ThermoFisher Scientific, J62688-AP), 0.5% NP-40 (Sigma-Aldrich, 74385), 1 mM DTT (ThermoFisher Scientific, Y00147), 200 U/mL RNase OUT (ThermoFisher Scientific, 10777019) and 1X EDTA-free Protease Inhibitor Cocktail (Roche, 11836170001). Then the lysate was incubated on ice for 5 minutes for thorough extraction and stored at −80 °C until immunoprecipitation. 50 μL of protein-G-magnetic beads (ThermoFisher Scientific, 10003D) were aliquoted into an RNase-free tube, placed on a magnetic rack and subjected to two washes with 500 μL NT-2 buffer containing 50 mM Tris-HCl, pH 7.4 (ThermoFisher Scientific, J60202-K2), 150 mM NaCl_2_ (Sigma-Aldrich, S9888), 1 mM MgCl_2_ (Sigma-Aldrich, M9272) and 0.05% NP-40 (Sigma-Aldrich, 74385). After removing the tube from the magnet, the beads were resuspended in 100 μL NT-2 buffer containing 5 μg of anti-IGF2BP2 antibody (Proteintech, 11601-1-AP). The resuspended beads were incubated by rotation for 1 hour at RT and spun and placed back on the magnetic rack. The supernatant was carefully aspirated, and the beads were thoroughly mixed and washed 6 times with 1 mL NT-2 buffer. Finally, the beads were resuspended in 900 μL NET-2 buffer containing 20 mM EDTA, pH 8.0 (Sigma-Aldrich, E9884), 1 mM DTT (ThermoFisher Scientific, Y00147), 200 U/mL RNase OUT (ThermoFisher Scientific, 10777019) supplemented with NT-2 -and kept on ice (referred below as ‘immunoprecipitation tube’). Additional 50 μL of magnetic beads were prepared simultaneously with 5 µg of anti-IgG (Proteintech, 30000-0-AP) for background assessment.

Lysates were thawed on ice and centrifuged at 20,000 g for 10 minutes at 4°C and 10 μL of the supernatant was collected as "Input" in an RNase-free tube, and stored at −80°C. 100 μL of the lysate supernatant was added to each immunoprecipitation tube, reaching a total reaction volume of 1000 μL. The tubes were then incubated on a rotator at 4°C overnight. Immunoprecipitation tubes were spun quickly and rinsed 6 times with 1 mL of pre-chilled NT-2 buffer. The beads-antibody-RNA binding protein complex was finally resuspended in 200 μL of cold NT-2 (referred to below as ‘RIP tube’). Simultaneously, NT-2 was added into the Input tube to achieve a final volume of 200 μL.

After the addition of 500 μL of TRI reagent (Sigma-Aldrich, T9424) and 100 µL of chloroform (Sigma-Aldrich, 650498), RIP tubes were vortexed vigorously and left for 5 minutes at RT. The tubes were then centrifuged at 16,000 g for 10 minutes at 4°C, and 300 μL of the aqueous phase were collected and transferred into a new RNase-free tube. To increase the RNA yield, 1X Precipitation Buffer (PB) containing 5 M ammonium acetate, 50 μL (Sigma-Aldrich, A7262), 7.5 M LiCl, 15 μL (Sigma-Aldrich, 310468); 25 μg glycogen (ThermoFisher Scientific, 10814010), 2-propanol, 600 μL (Sigma-Aldrich, 19516), was added and the tube placed at −80°C overnight. Tubes were centrifuged at 20,000 g for 30 minutes at 4 °C, and the pellet was washed twice with 500 μL 80% ethanol (Sigma-Aldrich, 32221-M) at 20,000 g for 10 minutes at 4°C. The supernatant was removed by aspiration and the pellet was air dried for 10-15minutes. Finally, the RNA pellet was resuspended with 20 μL of RNase-free water and stored at −80°C.

### Methylated (m^6^A) RNA immunoprecipitation (MeRIP)

Methylated (m^6^A) RNA immunoprecipitation (MeRIP) was performed according to a modification of a previous protocol (Dominissini et al., 2013). Briefly, the total RNA of 6 × 10^7^ M0 BMDMs at day 3 post-differentiation and *iv*LIMs at day 3 post-infection were extracted and the RNA concentration was adjusted to 1 µg/µL with RNase-free water (18 µL/tube). After saving 10% of fragmented RNA as “Input”, the remaining RNA was mixed with 200U RNasin (Promega, N2511), 200 µL of 5X Immunoprecipitation Buffer (IP) containing 50 mM Tris-HCl, pH 7.4, 750 mM NaCl_2_ and 0.5% NP-40 (Sigma-Aldrich, 74385) and 10 µg anti-m^6^A antibody (Proteintech, 68055-1-Ig) and adjusted with RNase-free water to 1 mL. A parallel reaction was set up with the same amount of fragmented RNA and 10 µg anti-IgG (Proteintech, 30000-0-AP) as a negative control. All immunoprecipitation samples were incubated on a rotator at 4°C overnight. Separately, 50 µL of Protein-G-Magnetic Beads (ThermoFisher Scientific, 10003D) were added to four RNase-free tubes, placed on the magnetic track, and washed three times with 1 mL of 1X IP buffer. Immunoprecipitation mixtures were then transferred to each tube containing pre-washed beads and incubated for 2 hours on a rotator at 4°C. Thereafter, the tubes were spun down, then placed on the pre-cold magnetic track and washed with 1 mL of 1X IP buffer five times. The supernatant was then discarded, and the bound RNA was resuspended with 200 µL of 1X IP buffer and extracted using 500 µL TRI reagent (Sigma-Aldrich, T9424). After the addition of 100 µL of chloroform (Sigma-Aldrich, 650498), tubes were incubated for 5 minutes at RT and centrifuged at 16,000 g for 10 minutes at 4°C. The aqueous phase was then carefully collected and transferred to a new RNase-free tube. Then PB-II containing 50 µL 5 M ammonium acetate (Sigma-Aldrich, A7262), 15 µL 7.5 M LiCl (Sigma-Aldrich, 310468), 25 μg glycogen (ThermoFisher Scientific, 10814010) and 600 µL 2-propanol (Sigma-Aldrich, 19516) was added to enhance the RNA precipitation overnight at −80°C. RNA was pelleted after centrifugation at 20,000 g for 30 minutes at 4°C and washed twice with 50 µL of 80% pre-cold ethanol (Sigma-Aldrich, 32221-M) at 20,000 g for 10 minutes at 4°C. The RNA pellet was air-dried for 10-15 minutes after discarding the supernatant and diluted in 20 µL RNase-free water for RT-qPCR.

### POSTAR3 analysis

POSTAR3, a public database for exploring RBPs binding site and associated downstream functional study, is based on publicly available large-scale cross-linking immunoprecipitation sequencing (CLIP-seq) datasets (Zhao et al., 2022).

## LEGENDS TO SUPPLEMENTARY FIGURES

**SUPPLEMENTARY FIGURE 1: Optimization and validation of M1 / M2 Mφ polarization conditions.**

M1 (A) and M2 (B) macrophage polarization states were induced by the addition of the indicated cocktails of cytokines for 72 hours to BMDMs at day 6 of *in vitro* differentiation. IFN-γ was added either at day 0 (D0) or day 2 (D2) of culture. Then Mφ viability (% YO-PRO-1 stained Mφs) and density (number of Hoechst 33342 stained Mφs per 0.3 mm2) were evaluated by epifluorescence microscopy analysis (EVOS system). The best M1 and 2 polarization conditions (indicated by purple and green histograms respectively) were selected and used for the entire study. (C) Transcriptomic validation of the selected polarization conditions. Total RNA was extracted and cDNA was produced for qPCR assays. The expression level of canonical M1 (upper panel) and M2 (lower panel) markers were analyzed in M0, M1 and M2 Mφs. Fold changes were calculated for M1 and M2 by using the M0 population as a calibrator. (D) Morphologic / phenotypic validation of the selected polarization conditions by combined brightfield and fluorescence microscopy analyses. Lipid droplets (LDs) were detected using the BODIPY 493/503 neutral lipid staining (green signal) and Mφ nuclei were stained by Hoechst 33342 (blue staining). Image acquisition was performed with the OPERA Phenix system. The exposition time for the BODIPY detection was defined on the M1 Mφs population and applied to all samples. LDs were only visualized in M1 macrophages. (E) Quantitation of secreted nitrites in Mφ supernatants. The supernatant of M0, M1, M2 Mφs and *iv*LIMs at day 3 PI were collected and secreted nitrites were quantified using the Griess assay. Bars represent the standard deviation of n = 2 independent biological replicates.

**SUPPLEMENTARY FIGURE 2: Characterization of the macrophage secretome.**

Culture supernatants of M0, M1, M2 macrophages and *iv*LIMs at day 3 post-infection were collected from three independent experiments. Supernatants corresponding to identical conditions were pooled for analysis. Samples were submitted to the Proteome Profiler Mouse Cytokine Array Kit to detect 111 analytes, *i.e*. cytokines, chemokines and acute phase proteins. Analytes were fixed on membranes through specific antibodies and were revealed by chemiluminescence. Imaging analysis was performed with the ImageQuantTL software. Pictures of the membranes corresponding to each group are shown (A). Examples of cytokine/chemokines secreted similarly by *iv*LIMs and M1, M2 or in an *iv*LIMs-specific way are indicated (CCL22, purple; TNF, green; CCL12, orange). Analyte/control locations are provided in the map (B) and the table (C) (light grey for Reference Spots; dark grey for Negative control).

**SUPPLEMENTARY FIGURE 3.**

**(A)** Fluorescence microscopy images of *iv*LIMs at days 3 and 30 PI and of *ev*LIMs isolated from lesions of infected RAG2 KO mice. Superpositions of fluorescence images obtained with Hoechst 33342 (nuclear staining, blue) and the mCherry signal of the transgenic parasites (red) are shown.

**(B)** Principal Coordinate Analysis (PCoA) of the expression values of a selection of transcripts in M1, M2 Mφs and *iv*LIMs. 58 transcripts for cytokines, chemokines, and members of the NF-κB pathway were assessed by RT-qPCR in M0, M1, M2 Mφs and *iv*LIMs at day 3 post-infection using the expression values of M0 Mφs as calibrators (n = 3 independent experiments). The numbers in the connecting lines indicate the distance between the three groups.

**SUPPLEMENTARY FIGURE 4. Simplified KEGG map representation of the pentose phosphate.**

(A) and fatty acid biosynthesis (B) pathways. Day 6 BMDMs were either infected with *L. amazonensis* amastigotes or left untreated. On day 3 PI total RNA was isolated and used for RNA-seq analysis. Significantly modulated genes in *iv*LIMs (adjusted p value < 0.05) are highlighted within the KEGG maps. Blue and red colors correspond to respectively down and up modulations observed in LIMs compared to uninfected M0 macrophages.

**SUPPLEMENTARY FIGURE 5. KEGG map of the fatty acid β oxydation pathway.**

Day 6 BMDMs were either infected with *L. amazonensis* amastigotes or left untreated. On day 3 PI total RNA was isolated and used for RNA-seq analysis. Significantly modulated genes in *iv*LIMs (adjusted p value < 0.05) are highlighted within the KEGG map. Blue and red colors correspond to respectively down and up modulations observed in LIMs compared to uninfected M0 macrophages.

**SUPPLEMENTARY FIGURE 6. Transcriptomic subversion of the electron transport chain in *iv*LIMs.**

Transcriptomic modulations were assessed by RNA-seq in day 3 *iv*LIMs *vs* M0 macrophages.

(A) Overall presentation of the Electron Transport Chain (ETC).

(B-C) Transcriptomic changes in *iv*LIMs of the different components of the NADH dehydrogenase (B), and the F-type ATPase and V-type ATPase (C) (blue: down modulation, red: up modulation).

**SUPPLEMENTARY FIGURE 7. Transcriptomic changes of the Tricarboxylic Acid Cycle pathway in *iv*LIMs.**

Transcriptomic modulations were assessed by RNA-seq in *iv*LIMs *vs* M0 macrophages and expression changes were projected onto a simplified KEGG map.

**SUPPLEMENTARY FIGURE 8. Adjustment of the protocol for the Seahorse XF Cell Mito stress kit for OCR analyses of *iv*LIMs.**

Oxygen Consumption Rates (OCR) of M0 macrophages, amastigotes and *iv*LIMs were measured with the Seahorse XF Cell Mito stress kit. (A) Presentation of the classical and modified protocols. Each protocol comprised three serial injections: 1: oligomycin (inhibitor of complex V); 2: FCCP (uncoupling agent of the mitochondrial electron transport chain), and 3: a mixture of rotenone (inhibitor of complex I) / antimycin A (inhibitor of complex III) (classical protocol); rotenone alone (modified protocol 1); antimycin A alone (modified protocol 2). (B) OCR determination during the application of these three protocols on M0 Mφs (1), isolated *L. amazonensis* amastigotes alone (2) and *iv*LIMs (3).

**SUPPLEMENTARY FIGURE 9. Cytoscape visualization of interaction map between RNA-Binding Proteins (RBPs) and glycolysis transcripts in *iv*LIMs.**

Data on RBPs and the targeted glycolysis transcripts were collected from the POSTAR3 public database. RNAs and RBPs are represented by hexagons and circles, respectively. The fold change of transcript abundance in *iv*LIMs compared to M0 BMDMs was extracted from the RNA-seq analysis. Genes that were up-regulated are colored in red, while down-regulated genes are colored in blue. The size of RBP circles indicates the number of target genes for selected glycolysis transcripts. Interactions between IGF2BP2 and its mRNA targets are highlighted in orange.

**SUPPLEMENTARY FIGURE 10. Evaluation of siRNA transfection efficiency in M0 BMDMs.**

Flow cytometric validation of the efficiency of our transfection procedure. Adherent M0 BMDMs were either untreated (1) or transfected for 3 hours without siRNA (1), with FITC-siRNA alone (2), with Transfection Reagent (TR) alone (3), or FITC-siRNA + TR (4). Then cells were carefully detached and analyzed by flow cytometry.

A) Biparametric dot plots displaying the fluorescence signals measured by the FITC and PerCP channels (x and y axis respectively). The percentage of macrophages located in the different quadrants are indicated.

B) Histograms of the fluorescence signals on the FITC channel.

## ACKNOWLEDGEMENTS

We acknowledge the financial support of the Institut Pasteur (Paris), the Région Ile-de-France (program DIM1Health), the Institut Pasteur International Direction to the International Mixed Unit “Inflammation and *Leishmania* Infection”, the ERC Synergy project DecoLeishRN (Grant agreement ID: 101071613), and the Institut National de la santé et de la recherche médicale (INSERM). Sheng Zhang is part of the Pasteur - Paris University (PPU) International PhD program.

## Notes

### Competing Interest Statement

The authors have declared no competing interest.

### Summary of Updates

This new version updates and completes our first manuscript submited in 2022. It provides new mechanistic insight into a potential epi-transcriptomic process underlying increased glycolytic activity in macrophages infected by Leishmania amazonensis amastigotes (LIMs). New figure 1 combines previous figures 1 and 2. New figure 2 replaces previous figures 3 and 4 by providing a more precise metabolic analyses of LIMs using an adapted protocol of the Seahorse assays that now takes the parasite activity into account. It also included previous supplementary figure 7 (S7). New figures 3 and 4 propose new data that explain how glycolysis is induced in LIMs. New figure 3 shows that the increased glycolysis of LIMs is not associated to HIF-1alpha up regulation but to increased expression of the m6A reader protein IGF2BP2 and its effect on stabilizing Hk2 mRNA via its increased m6A modification. New Figure 4 shows that IGF2BP2 knockdown reduces glycolytic activity in LIMs, thus genetically confirming its role in regulating energy metabolism in LIMs. New Figure 5 shows a model that updates and summerizes all new data. Supplementary figures were also modified. New S1 combines previous S1 and S3. Previous S2 was removed being redundant with new S1. New S2 corresponds to previous S4. New S3 corresponds to an updated version of previous figure 1C and previous figure 1E. New S4 and S5 present KEGG analyses of the PPP projecting RNAseq expression data of LIMs versus non-infected macrophages (previous S8). All the following supplementary figures show new data. New S4 and S5 present KEGG analyses of fatty acid synthesis and oxidation projecting RNAseq expression data of LIMs versus non-infected macrophages. New S6 and S7 present KEGG analyses of the oxydative phoshorylation and the TCA cycle and include previous S9. New S8 presents the adaptations of the Seahorse assay. New S9 presents RNA-binding protein analyses. New S10 presents flow cytometry analysis of siRNA transfection efficiency in M0 macrophages.

