## Supplemental table 1 for "The m^6^A reader IGF2BP2 directs immune-metabolic reprogramming in Leishmania amazonensis-infected macrophages"

### M1 list

| Gene name | ENSEMBL Id | Description |
|---|---|---|
| Abcc5 | ENSMUSG00000022822 | ATP-binding cassette, sub-family C (CFTR/MRP), member 5 |
| Acod1 | ENSMUSG00000022126 | aconitate decarboxylase 1 |
| Acp2 | ENSMUSG00000002103 | acid phosphatase 2, lysosomal |
| Acp5 | ENSMUSG00000001348 | acid phosphatase 5, tartrate resistant |
| Acsl1 | ENSMUSG00000018796 | acyl-CoA synthetase long-chain family member 1 |
| Adgb | ENSMUSG00000050994 | androglobin |
| Adora2a | ENSMUSG00000020178 | adenosine A2a receptor |
| Aif1 | ENSMUSG00000024397 | allograft inflammatory factor 1 |
| Ankrd37 | ENSMUSG00000050914 | ankyrin repeat domain 37 |
| Antxr1 | ENSMUSG00000033420 | anthrax toxin receptor 1 |
| Aoah | ENSMUSG00000021322 | acyloxyacyl hydrolase |
| Arg2 | ENSMUSG00000021125 | arginase type II |
| Batf | ENSMUSG00000034266 | basic leucine zipper transcription factor, ATF-like |
| Bcl3 | ENSMUSG00000053175 | B cell leukemia/lymphoma 3 |
| Birc3 | ENSMUSG00000032000 | baculoviral IAP repeat-containing 3 |
| Bnip3 | ENSMUSG00000078566 | BCL2/adenovirus E1B interacting protein 3 |
| Bst1 | ENSMUSG00000029082 | bone marrow stromal cell antigen 1 |
| Car13 | ENSMUSG00000027555 | carbonic anhydrase 13 |
| Carmil1 | ENSMUSG00000021338 | capping protein regulator and myosin 1 linker 1 |
| Ccl5 | ENSMUSG00000035042 | chemokine (C-C motif) ligand 5 |
| Ccrl2 | ENSMUSG00000043953 | chemokine (C-C motif) receptor-like 2 |
| Cd200 | ENSMUSG00000022661 | CD200 molecule |
| Cd274 | ENSMUSG00000016496 | CD274 antigen |
| Cd38 | ENSMUSG00000029084 | CD38 antigen |
| Cd40 | ENSMUSG00000017652 | CD40 antigen |
| Cd69 | ENSMUSG00000030156 | CD69 antigen |
| Cd86 | ENSMUSG00000022901 | CD86 antigen |
| Cdc42ep2 | ENSMUSG00000045664 | CDC42 effector protein (Rho GTPase binding) 2 |
| Cebpb | ENSMUSG00000056501 | CCAAT/enhancer binding protein (C/EBP), beta |
| Cflar | ENSMUSG00000026031 | CASP8 and FADD-like apoptosis regulator |
| Clec4e | ENSMUSG00000030142 | C-type lectin domain family 4, member e |
| Clmp | ENSMUSG00000032024 | CXADR-like membrane protein |
| Cp | ENSMUSG00000003617 | ceruloplasmin |
| Cxcl16 | ENSMUSG00000018920 | chemokine (C-X-C motif) ligand 16 |
| Cxcl2 | ENSMUSG00000058427 | chemokine (C-X-C motif) ligand 2 |
| Cxcl3 | ENSMUSG00000029379 | chemokine (C-X-C motif) ligand 3 |
| Cxcl9 | ENSMUSG00000029417 | chemokine (C-X-C motif) ligand 9 |
| Cybb | ENSMUSG00000015340 | cytochrome b-245, beta polypeptide |
| Ddx60 | ENSMUSG00000037921 | DExD/H box helicase 60 |
| Dnmt3l | ENSMUSG00000000730 | DNA (cytosine-5-)-methyltransferase 3-like |
| Dram1 | ENSMUSG00000020057 | DNA-damage regulated autophagy modulator 1 |
| Dst | ENSMUSG00000026131 | dystonin |
| Dtx2 | ENSMUSG00000004947 | deltex 2, E3 ubiquitin ligase |
| Dusp1 | ENSMUSG00000024190 | dual specificity phosphatase 1 |
| Ebi3 | ENSMUSG00000003206 | Epstein-Barr virus induced gene 3 |
| Egln3 | ENSMUSG00000035105 | egl-9 family hypoxia-inducible factor 3 |
| Ehd1 | ENSMUSG00000024772 | EH-domain containing 1 |
| Ell2 | ENSMUSG00000001542 | elongation factor for RNA polymerase II 2 |
| Epb41l3 | ENSMUSG00000024044 | erythrocyte membrane protein band 4.1 like 3 |
| Ero1a | ENSMUSG00000021831 | endoplasmic reticulum oxidoreductase 1 alpha |
| Eva1b | ENSMUSG00000050212 | eva-1 homolog B (C. elegans) |
| Fam114a1 | ENSMUSG00000029185 | family with sequence similarity 114, member A1 |
| Ficd | ENSMUSG00000053334 | FIC domain containing |
| Fpr1 | ENSMUSG00000045551 | formyl peptide receptor 1 |
| Gadd45b | ENSMUSG00000015312 | growth arrest and DNA-damage-inducible 45 beta |
| Gch1 | ENSMUSG00000037580 | GTP cyclohydrolase 1 |
| Gcnt2 | ENSMUSG00000021360 | glucosaminyl (N-acetyl) transferase 2, I-branching enzyme |
| Gde1 | ENSMUSG00000033917 | glycerophosphodiester phosphodiesterase 1 |
| Ggct | ENSMUSG00000002797 | gamma-glutamyl cyclotransferase |
| Gngt2 | ENSMUSG00000038811 | guanine nucleotide binding protein (G protein), gamma transducing activity polypeptide 2 |
| Gpd2 | ENSMUSG00000026827 | glycerol phosphate dehydrogenase 2, mitochondrial |
| Gsap | ENSMUSG00000039934 | gamma-secretase activating protein |
| Gsr | ENSMUSG00000031584 | glutathione reductase |
| Gstt1 | ENSMUSG00000001663 | glutathione S-transferase, theta 1 |

| Gene name | ENSEMBL Id | Description |
|---|---|---|
| H2-Ob | ENSMUSG00000041538 | histocompatibility 2, O region beta locus |
| H2-Q5 | ENSMUSG00000055413 | histocompatibility 2, Q region locus 5 |
| Hif1a | ENSMUSG00000021109 | hypoxia inducible factor 1, alpha subunit |
| Hmox1 | ENSMUSG00000005413 | heme oxygenase 1 |
| Hp | ENSMUSG00000031722 | haptoglobin |
| Hspa1a | ENSMUSG00000091971 | heat shock protein 1A |
| Hspa1b | ENSMUSG00000090877 | heat shock protein 1B |
| Id3 | ENSMUSG00000007872 | inhibitor of DNA binding 3 |
| Ifi209 | ENSMUSG00000043263 | interferon activated gene 209 |
| Ifi27l2a | ENSMUSG00000079017 | interferon, alpha-inducible protein 27 like 2A |
| Ifi44 | ENSMUSG00000028037 | interferon-induced protein 44 |
| Ifit2 | ENSMUSG00000045932 | interferon-induced protein with tetratricopeptide repeats 2 |
| Ifitm6 | ENSMUSG00000059108 | interferon induced transmembrane protein 6 |
| Ifnar2 | ENSMUSG00000022971 | interferon (alpha and beta) receptor 2 |
| Il15ra | ENSMUSG00000023206 | interleukin 15 receptor, alpha chain |
| Il1a | ENSMUSG00000027399 | interleukin 1 alpha |
| Il1b | ENSMUSG00000027398 | interleukin 1 beta |
| Il1rn | ENSMUSG00000026981 | interleukin 1 receptor antagonist |
| Irak2 | ENSMUSG00000060477 | interleukin-1 receptor-associated kinase 2 |
| Irak3 | ENSMUSG00000020227 | interleukin-1 receptor-associated kinase 3 |
| Isg20 | ENSMUSG00000039236 | interferon-stimulated protein |
| Itga4 | ENSMUSG00000027009 | integrin alpha 4 |
| Itga5 | ENSMUSG00000000555 | integrin alpha 5 (fibronectin receptor alpha) |
| Itgal | ENSMUSG00000030830 | integrin alpha L |
| Jag1 | ENSMUSG00000027276 | jagged 1 |
| Klra2 | ENSMUSG00000030187 | killer cell lectin-like receptor, subfamily A, member 2 |
| Lcn2 | ENSMUSG00000026822 | lipocalin 2 |
| Lox | ENSMUSG00000024529 | lysyl oxidase |
| Marchf1 | ENSMUSG00000036469 | membrane associated ring-CH-type finger 1 |
| Marco | ENSMUSG00000026390 | macrophage receptor with collagenous structure |
| Mcemp1 | ENSMUSG00000013974 | mast cell expressed membrane protein 1 |
| Mdm2 | ENSMUSG00000020184 | transformed mouse 3T3 cell double minute 2 |
| Mefv | ENSMUSG00000022534 | Mediterranean fever |
| Met | ENSMUSG00000009376 | met proto-oncogene |
| Mfap3l | ENSMUSG00000031647 | microfibrillar-associated protein 3-like |
| Mlkl | ENSMUSG00000012519 | mixed lineage kinase domain-like |
| Mmp14 | ENSMUSG00000000957 | matrix metallopeptidase 14 (membrane-inserted) |
| Ms4a4c | ENSMUSG00000024675 | membrane-spanning 4-domains, subfamily A, member 4C |
| Ms4a6d | ENSMUSG00000024679 | membrane-spanning 4-domains, subfamily A, member 6D |
| Mx1 | ENSMUSG00000000386 | MX dynamin-like GTPase 1 |
| Mxd1 | ENSMUSG00000001156 | MAX dimerization protein 1 |
| Ndufaf3 | ENSMUSG00000070283 | NADH:ubiquinone oxidoreductase complex assembly factor 3 |
| Nfkbia | ENSMUSG00000021025 | nuclear factor of kappa light polypeptide gene enhancer in B cells inhibitor, alpha |
| Nfkbiz | ENSMUSG00000035356 | nuclear factor of kappa light polypeptide gene enhancer in B cells inhibitor, zeta |
| Nos2 | ENSMUSG00000020826 | nitric oxide synthase 2, inducible |
| Nupr1 | ENSMUSG00000030717 | nuclear protein transcription regulator 1 |
| Oasl1 | ENSMUSG00000041827 | 2'-5' oligoadenylate synthetase-like 1 |
| P4ha1 | ENSMUSG00000019916 | procollagen-proline, 2-oxoglutarate 4-dioxygenase (proline 4-hydroxylase), alpha 1 polypeptide |
| Pdpn | ENSMUSG00000028583 | podoplanin |
| Pfkfb3 | ENSMUSG00000026773 | 6-phosphofructo-2-kinase/fructose-2,6-biphosphatase 3 |
| Pid1 | ENSMUSG00000045658 | phosphotyrosine interaction domain containing 1 |
| Pilra | ENSMUSG00000046245 | paired immunoglobin-like type 2 receptor alpha |
| Plpp1 | ENSMUSG00000021759 | phospholipid phosphatase 1 |
| Plpp3 | ENSMUSG00000028517 | phospholipid phosphatase 3 |
| Ppp1r12b | ENSMUSG00000073557 | protein phosphatase 1, regulatory subunit 12B |
| Prdx5 | ENSMUSG00000024953 | peroxiredoxin 5 |
| Procr | ENSMUSG00000027611 | protein C receptor, endothelial |
| Prss46 | ENSMUSG00000049719 | protease, serine 46 |
| Psmd10 | ENSMUSG00000031429 | proteasome (prosome, macropain) 26S subunit, non-ATPase, 10 |
| Pstpip2 | ENSMUSG00000025429 | proline-serine-threonine phosphatase-interacting protein 2 |
| Ptges | ENSMUSG00000050737 | prostaglandin E synthase |
| Ptgs2 | ENSMUSG00000032487 | prostaglandin-endoperoxide synthase 2 |
| Pvr | ENSMUSG00000040511 | poliovirus receptor |
| Rab20 | ENSMUSG00000031504 | RAB20, member RAS oncogene family |
| Rbpms | ENSMUSG00000031586 | RNA binding protein gene with multiple splicing |
| Rffl | ENSMUSG00000020696 | ring finger and FYVE like domain containing protein |

| Gene name | ENSEMBL Id | Description |
|---|---|---|
| Rims3 | ENSMUSG00000032890 | regulating synaptic membrane exocytosis 3 |
| Rnd1 | ENSMUSG00000054855 | Rho family GTPase 1 |
| Saa3 | ENSMUSG00000040026 | serum amyloid A 3 |
| Samsn1 | ENSMUSG00000022876 | SAM domain, SH3 domain and nuclear localization signals, 1 |
| Sdc1 | ENSMUSG00000020592 | syndecan 1 |
| Serf1 | ENSMUSG00000021643 | small EDRK-rich factor 1 |
| Serpine1 | ENSMUSG00000037411 | serine (or cysteine) peptidase inhibitor, clade E, member 1 |
| Sgms2 | ENSMUSG00000050931 | sphingomyelin synthase 2 |
| Slamf6 | ENSMUSG00000015314 | SLAM family member 6 |
| Slamf7 | ENSMUSG00000038179 | SLAM family member 7 |
| Slc15a3 | ENSMUSG00000024737 | solute carrier family 15, member 3 |
| Slc16a3 | ENSMUSG00000025161 | solute carrier family 16 (monocarboxylic acid transporters), member 3 |
| Slc22a4 | ENSMUSG00000020334 | solute carrier family 22 (organic cation transporter), member 4 |
| Slc25a33 | ENSMUSG00000028982 | solute carrier family 25, member 33 |
| Slc31a2 | ENSMUSG00000066152 | solute carrier family 31, member 2 |
| Slc43a3 | ENSMUSG00000027074 | solute carrier family 43, member 3 |
| Slc6a12 | ENSMUSG00000030109 | solute carrier family 6 (neurotransmitter transporter, betaine/GABA), member 12 |
| Slc7a8 | ENSMUSG00000022180 | solute carrier family 7 (cationic amino acid transporter, y+ system), member 8 |
| Slco3a1 | ENSMUSG00000025790 | solute carrier organic anion transporter family, member 3a1 |
| Slfn1 | ENSMUSG00000078763 | schlafen 1 |
| Slfn4 | ENSMUSG00000000204 | schlafen 4 |
| Slpi | ENSMUSG00000017002 | secretory leukocyte peptidase inhibitor |
| Smad6 | ENSMUSG00000036867 | SMAD family member 6 |
| Smim3 | ENSMUSG00000038059 | small integral membrane protein 3 |
| Smpdl3b | ENSMUSG00000028885 | sphingomyelin phosphodiesterase, acid-like 3B |
| Snapc1 | ENSMUSG00000021113 | small nuclear RNA activating complex, polypeptide 1 |
| Snx18 | ENSMUSG00000042364 | sorting nexin 18 |
| Socs3 | ENSMUSG00000053113 | suppressor of cytokine signaling 3 |
| Sod2 | ENSMUSG00000006818 | superoxide dismutase 2, mitochondrial |
| Spic | ENSMUSG00000004359 | Spi-C transcription factor (Spi-1/PU.1 related) |
| St3gal5 | ENSMUSG00000056091 | ST3 beta-galactoside alpha-2,3-sialyltransferase 5 |
| Stx11 | ENSMUSG00000039232 | syntaxin 11 |
| Susd2 | ENSMUSG00000006342 | sushi domain containing 2 |
| Tapt1 | ENSMUSG00000046985 | transmembrane anterior posterior transformation 1 |
| Thbs1 | ENSMUSG00000040152 | thrombospondin 1 |
| Tlr2 | ENSMUSG00000027995 | toll-like receptor 2 |
| Tma16 | ENSMUSG00000025591 | translation machinery associated 16 |
| Tmem119 | ENSMUSG00000054675 | transmembrane protein 119 |
| Tnf | ENSMUSG00000024401 | tumor necrosis factor |
| Tnfaip3 | ENSMUSG00000019850 | tumor necrosis factor, alpha-induced protein 3 |
| Tnfrsf1b | ENSMUSG00000028599 | tumor necrosis factor receptor superfamily, member 1b |
| Tnfrsf21 | ENSMUSG00000023915 | tumor necrosis factor receptor superfamily, member 21 |
| Tnfsf9 | ENSMUSG00000035678 | tumor necrosis factor (ligand) superfamily, member 9 |
| Traf1 | ENSMUSG00000026875 | TNF receptor-associated factor 1 |
| Trem3 | ENSMUSG00000041754 | triggering receptor expressed on myeloid cells 3 |
| Tspan3 | ENSMUSG00000032324 | tetraspanin 3 |
| Tspo | ENSMUSG00000041736 | translocator protein |
| Tspoap1 | ENSMUSG00000034156 | TSPO associated protein 1 |
| Usp18 | ENSMUSG00000030107 | ubiquitin specific peptidase 18 |
| Vcam1 | ENSMUSG00000027962 | vascular cell adhesion molecule 1 |
| Vcan | ENSMUSG00000021614 | versican |
| Vegfa | ENSMUSG00000023951 | vascular endothelial growth factor A |
| Wfdc18 | ENSMUSG00000000983 | WAP four-disulfide core domain 18 |
| Wnt6 | ENSMUSG00000033227 | wingless-type MMTV integration site family, member 6 |
| Xaf1 | ENSMUSG00000040483 | XIAP associated factor 1 |
| Zbp1 | ENSMUSG00000027514 | Z-DNA binding protein 1 |
| Zbtb7b | ENSMUSG00000028042 | zinc finger and BTB domain containing 7B |
| Zfp503 | ENSMUSG00000039081 | zinc finger protein 503 |
| Zfp658 | ENSMUSG00000056592 | zinc finger protein 658 |
