## Supplemental table 2 for "The m^6^A reader IGF2BP2 directs immune-metabolic reprogramming in Leishmania amazonensis-infected macrophages"

### M2 list

| Gene name | ENSEMBL Id | Description |
|---|---|---|
| Acap1 | ENSMUSG00000001588 | ArfGAP with coiled-coil, ankyrin repeat and PH domains 1 |
| Adipoq | ENSMUSG00000022878 | adiponectin, C1Q and collagen domain containing |
| Ahnak | ENSMUSG00000069833 | AHNAK nucleoprotein (desmoyokin) |
| Ahr | ENSMUSG00000019256 | aryl-hydrocarbon receptor |
| Alms1 | ENSMUSG00000063810 | ALMS1, centrosome and basal body associated |
| Ap2m1 | ENSMUSG00000022841 | adaptor-related protein complex 2, mu 1 subunit |
| Arg1 | ENSMUSG00000019987 | arginase, liver |
| Arhgap6 | ENSMUSG00000031355 | Rho GTPase activating protein 6 |
| Arl15 | ENSMUSG00000042348 | ADP-ribosylation factor-like 15 |
| Atp6v0a1 | ENSMUSG00000019302 | ATPase, H+ transporting, lysosomal V0 subunit A1 |
| Atp6v0d2 | ENSMUSG00000028238 | ATPase, H+ transporting, lysosomal V0 subunit D2 |
| Batf3 | ENSMUSG00000026630 | basic leucine zipper transcription factor, ATF-like 3 |
| Bcar3 | ENSMUSG00000028121 | breast cancer anti-estrogen resistance 3 |
| Bhlhe40 | ENSMUSG00000030103 | basic helix-loop-helix family, member e40 |
| Car2 | ENSMUSG00000027562 | carbonic anhydrase 2 |
| Car5b | ENSMUSG00000031373 | carbonic anhydrase 5b, mitochondrial |
| Casp6 | ENSMUSG00000027997 | caspase 6 |
| Cblb | ENSMUSG00000022637 | Casitas B-lineage lymphoma b |
| Ccdc88a | ENSMUSG00000032740 | coiled coil domain containing 88A |
| Ccl17 | ENSMUSG00000031780 | chemokine (C-C motif) ligand 17 |
| Ccl22 | ENSMUSG00000031779 | chemokine (C-C motif) ligand 22 |
| Ccl24 | ENSMUSG00000004814 | chemokine (C-C motif) ligand 24 |
| Cd300ld | ENSMUSG00000034641 | CD300 molecule like family member d |
| Cd74 | ENSMUSG00000024610 | CD74 antigen (invariant polypeptide of major histocompatibility complex, class II antigen-associated) |
| Ch25h | ENSMUSG00000050370 | cholesterol 25-hydroxylase |
| Chil3 | ENSMUSG00000040809 | chitinase-like 3 |
| Chst11 | ENSMUSG00000034612 | carbohydrate sulfotransferase 11 |
| Ciita | ENSMUSG00000022504 | class II transactivator |
| Cish | ENSMUSG00000032578 | cytokine inducible SH2-containing protein |
| Clec10a | ENSMUSG00000000318 | C-type lectin domain family 10, member A |
| Clec7a | ENSMUSG00000079293 | C-type lectin domain family 7, member a |
| Clic4 | ENSMUSG00000037242 | chloride intracellular channel 4 (mitochondrial) |
| Cttn | ENSMUSG00000031078 | cortactin |
| Cyp1b1 | ENSMUSG00000024087 | cytochrome P450, family 1, subfamily b, polypeptide 1 |
| Cytip | ENSMUSG00000026832 | cytohesin 1 interacting protein |
| Dclk1 | ENSMUSG00000027797 | doublecortin-like kinase 1 |
| Dcstamp | ENSMUSG00000022303 | dendrocyte expressed seven transmembrane protein |
| Ddhd1 | ENSMUSG00000037697 | DDHD domain containing 1 |
| Diras2 | ENSMUSG00000047842 | DIRAS family, GTP-binding RAS-like 2 |
| Dlg3 | ENSMUSG00000000881 | discs large MAGUK scaffold protein 3 |
| Dusp4 | ENSMUSG00000031530 | dual specificity phosphatase 4 |
| Edn1 | ENSMUSG00000021367 | endothelin 1 |
| Egr2 | ENSMUSG00000037868 | early growth response 2 |
| Emp2 | ENSMUSG00000022505 | epithelial membrane protein 2 |
| Fam20c | ENSMUSG00000025854 | FAM20C, golgi associated secretory pathway kinase |
| Fchsd2 | ENSMUSG00000030691 | FCH and double SH3 domains 2 |
| Ffar4 | ENSMUSG00000054200 | free fatty acid receptor 4 |
| Flrt2 | ENSMUSG00000047414 | fibronectin leucine rich transmembrane protein 2 |
| Flt1 | ENSMUSG00000029648 | FMS-like tyrosine kinase 1 |
| Fn1 | ENSMUSG00000026193 | fibronectin 1 |
| Fyn | ENSMUSG00000019843 | Fyn proto-oncogene |
| Gab1 | ENSMUSG00000031714 | growth factor receptor bound protein 2-associated protein 1 |
| Gas6 | ENSMUSG00000031451 | growth arrest specific 6 |
| Gask1b | ENSMUSG00000027955 | golgi associated kinase 1B |
| Gda | ENSMUSG00000058624 | guanine deaminase |
| Gnb4 | ENSMUSG00000027669 | guanine nucleotide binding protein (G protein), beta 4 |
| Gpc1 | ENSMUSG00000034220 | glypican 1 |
| Gpr155 | ENSMUSG00000041762 | G protein-coupled receptor 155 |
| H2-Aa | ENSMUSG00000036594 | histocompatibility 2, class II antigen A, alpha |
| H2-Eb1 | ENSMUSG00000060586 | histocompatibility 2, class II antigen E beta |
| Hacd1 | ENSMUSG00000063275 | 3-hydroxyacyl-CoA dehydratase 1 |
| Hbegf | ENSMUSG00000024486 | heparin-binding EGF-like growth factor |
| Hebp2 | ENSMUSG00000019853 | heme binding protein 2 |
| Iars | ENSMUSG00000037851 | isoleucine-tRNA synthetase |
| Il1rl1 | ENSMUSG00000026069 | interleukin 1 receptor-like 1 |
| Il1rl2 | ENSMUSG00000070942 | interleukin 1 receptor-like 2 |

| Gene name | ENSEMBL Id | Description |
|---|---|---|
| Il27ra | ENSMUSG00000005465 | interleukin 27 receptor, alpha |
| Il31ra | ENSMUSG00000050377 | interleukin 31 receptor A |
| Il6st | ENSMUSG00000021756 | interleukin 6 signal transducer |
| Inpp5a | ENSMUSG00000025477 | inositol polyphosphate-5-phosphatase A |
| Irf4 | ENSMUSG00000021356 | interferon regulatory factor 4 |
| Itga1 | ENSMUSG00000042284 | integrin alpha 1 |
| Itgax | ENSMUSG00000030789 | integrin alpha X |
| Itgb3 | ENSMUSG00000020689 | integrin beta 3 |
| Klf4 | ENSMUSG00000003032 | Kruppel-like transcription factor 4 (gut) |
| Klf9 | ENSMUSG00000033863 | Kruppel-like transcription factor 9 |
| Limk1 | ENSMUSG00000029674 | LIM domain kinase 1 |
| Map4k1 | ENSMUSG00000037337 | mitogen-activated protein kinase kinase kinase kinase 1 |
| Matk | ENSMUSG00000004933 | megakaryocyte-associated tyrosine kinase |
| Mgl2 | ENSMUSG00000040950 | macrophage galactose N-acetyl-galactosamine specific lectin 2 |
| Mmp12 | ENSMUSG00000049723 | matrix metallopeptidase 12 |
| Mmp19 | ENSMUSG00000025355 | matrix metallopeptidase 19 |
| Mrc1 | ENSMUSG00000026712 | mannose receptor, C type 1 |
| Myc | ENSMUSG00000022346 | myelocytomatosis oncogene |
| Myrf | ENSMUSG00000036098 | myelin regulatory factor |
| Nek6 | ENSMUSG00000026749 | NIMA (never in mitosis gene a)-related expressed kinase 6 |
| Niban2 | ENSMUSG00000026796 | niban apoptosis regulator 2 |
| Ocstamp | ENSMUSG00000027670 | osteoclast stimulatory transmembrane protein |
| Olr1 | ENSMUSG00000030162 | oxidized low density lipoprotein (lectin-like) receptor 1 |
| P2ry1 | ENSMUSG00000027765 | purinergic receptor P2Y, G-protein coupled 1 |
| Papss2 | ENSMUSG00000024899 | 3'-phosphoadenosine 5'-phosphosulfate synthase 2 |
| Pdcd1lg2 | ENSMUSG00000016498 | programmed cell death 1 ligand 2 |
| Pde12 | ENSMUSG00000043702 | phosphodiesterase 12 |
| Pdia4 | ENSMUSG00000025823 | protein disulfide isomerase associated 4 |
| Pex26 | ENSMUSG00000067825 | peroxisomal biogenesis factor 26 |
| Pfkp | ENSMUSG00000021196 | phosphofructokinase, platelet |
| Pkp2 | ENSMUSG00000041957 | plakophilin 2 |
| Plbd1 | ENSMUSG00000030214 | phospholipase B domain containing 1 |
| Plekhf1 | ENSMUSG00000074170 | pleckstrin homology domain containing, family F (with FYVE domain) member 1 |
| Plk2 | ENSMUSG00000021701 | polo like kinase 2 |
| Plxdc2 | ENSMUSG00000026748 | plexin domain containing 2 |
| Plxnc1 | ENSMUSG00000074785 | plexin C1 |
| Prkar1b | ENSMUSG00000025855 | protein kinase, cAMP dependent regulatory, type I beta |
| Ptgs1 | ENSMUSG00000047250 | prostaglandin-endoperoxide synthase 1 |
| Ptpre | ENSMUSG00000041836 | protein tyrosine phosphatase receptor type E |
| Ralgds | ENSMUSG00000026821 | ral guanine nucleotide dissociation stimulator |
| Rassf8 | ENSMUSG00000030259 | Ras association (RalGDS/AF-6) domain family (N-terminal) member 8 |
| Rgs16 | ENSMUSG00000026475 | regulator of G-protein signaling 16 |
| Rhoj | ENSMUSG00000046768 | ras homolog family member J |
| Rnase2a | ENSMUSG00000047222 | ribonuclease, RNase A family, 2A (liver, eosinophil-derived neurotoxin) |
| Rras2 | ENSMUSG00000055723 | related RAS viral (r-ras) oncogene 2 |
| Sema4b | ENSMUSG00000030539 | sema domain, immunoglobulin domain (Ig), transmembrane domain (TM) and short cytoplasmic domain, (semaphorin) 4B |
| Sema6d | ENSMUSG00000027200 | sema domain, transmembrane domain (TM), and cytoplasmic domain, (semaphorin) 6D |
| Septin11 | ENSMUSG00000058013 | septin 11 |
| Siglecf | ENSMUSG00000039013 | sialic acid binding Ig-like lectin F |
| Slc25a13 | ENSMUSG00000015112 | solute carrier family 25 (mitochondrial carrier, adenine nucleotide translocator), member 13 |
| Slc9a9 | ENSMUSG00000031129 | solute carrier family 9 (sodium/hydrogen exchanger), member 9 |
| Smagp | ENSMUSG00000053559 | small cell adhesion glycoprotein |
| Smap2 | ENSMUSG00000032870 | small ArfGAP 2 |
| Snn | ENSMUSG00000037972 | stannin |
| Socs2 | ENSMUSG00000020027 | suppressor of cytokine signaling 2 |
| Socs6 | ENSMUSG00000056153 | suppressor of cytokine signaling 6 |
| St8sia1 | ENSMUSG00000030283 | ST8 alpha-N-acetyl-neuraminide alpha-2,8-sialyltransferase 1 |
| Tanc2 | ENSMUSG00000053580 | tetratricopeptide repeat, ankyrin repeat and coiled-coil containing 2 |
| Tbc1d16 | ENSMUSG00000039976 | TBC1 domain family, member 16 |
| Tfrc | ENSMUSG00000022797 | transferrin receptor |
| Tiam1 | ENSMUSG00000002489 | T cell lymphoma invasion and metastasis 1 |
| Tmem26 | ENSMUSG00000060044 | transmembrane protein 26 |
| Traf5 | ENSMUSG00000026637 | TNF receptor-associated factor 5 |
| Tspan5 | ENSMUSG00000028152 | tetraspanin 5 |
| Tuba8 | ENSMUSG00000030137 | tubulin, alpha 8 |
| Vwf | ENSMUSG00000001930 | Von Willebrand factor |
| Wdfy2 | ENSMUSG00000014547 | WD repeat and FYVE domain containing 2 |
| Zyx | ENSMUSG00000029860 | Zyxin |
