## Supplemental table 3 for "The m^6^A reader IGF2BP2 directs immune-metabolic reprogramming in Leishmania amazonensis-infected macrophages"

**Zhang et al., Supplementary Table 3**

| Transcript | Forward | Reverse |
|---|---|---|
| ***Aldoa*** | 5'-GGAACCAATGGCGAGACAA-3' | 5'-TCGGCTCCATCCTTCTTATAC-3' |
| ***Arg1*** | 5'-CTCCAAGCCAAAGTCCTTAGAG-3' | 5'-AGGAGCTGTCATTAGGGACATC-3' |
| ***Btk*** | 5'-ATGTAATCCGGTACAATAGTGAC-3' | 5'-GCTTCCATTCCTGTTCTCC-3' |
| ***Ccl1*** | 5'-AGTGTTACAGAAAGATGGGC-3' | 5'-TTGTTAGTTGAGGCGCAG-3' |
| ***Ccl17*** | 5'-GCCATTCCTATCAGGAAGTTG-3' | 5'-CTGGACAGTCAGAAACACG-3' |
| ***Ccl2*** | 5'-GTCCCAAAGAAGCTGTAGT-3' | 5'-TATGTCTGGACCCATTCCT-3' |
| ***Ccl3*** | 5'-TCCACGCCAATTCATCG-3' | 5'-ATCTGCCGGTTTCTCTTAG-3' |
| ***Ccr2*** | 5'-CACGGCATACTATCAACATCT-3' | 5'-CAATTTGCTTCACACTGGTTT-3' |
| ***Cd68*** | 5'-TGTCTGATCTTGCTAGGACCG-3' | 5'-GAGAGTAACGGCCTTTTTGTGA-3' |
| ***Cd80*** | 5'-ATACACCACTCCTCAAGTTT-3' | 5'-ACTGACTTGGACAGTTGTTC-3' |
| ***Cd86*** | 5'-CACAATGTTCAGATCAAGGAC-3' | 5'-GTTCACTGAAGTTGGCGA-3' |
| ***Cebpd*** | 5'-TGTGCCACGACGAACTC-3' | 5'-CCTAGCGACAGACCCCA-3' |
| ***Chil1*** | 5'-CGAGATATGCGACTTCCTCA-3' | 5'-CACTGGTTGCCCTTGGTA-3' |
| ***Chuk*** | 5'- CTGGGATCCGATATCTGC -3' | 5'- TGGCTGTGTACGGCTTA -3' |
| ***Cxcl1*** | 5'-CTGGGATTCACCTCAAGAAC-3' | 5'-GAGTGTGGCTATGACTTCG-3' |
| ***Cxcl10*** | 5'-CAGTGAGAATGAGGGCCATA-3' | 5'-ATCGTGGCAATGATCTCAACA-3' |
| ***Cxcl11*** | 5'-CTGCGACAAAGTTGAAGTGA-3' | 5'-ATTATGAGGCGAGCTTGC-3' |
| ***Cxcl9*** | 5'-GTTCGAGGAACCCTAGTGA-3' | 5'-GTCTTTGAGGGATTTGTAGTGG-3' |
| ***Dhcr24*** | 5'-ACGTGTGAAGCACTTCG-3' | 5'-CTCCCAGAATTCCTCGC-3' |
| ***Eno1*** | 5'-GCACTCAGAAAGTGAATGTTG-3' | 5'-AGATTTATTCTCTGTGCCGTC-3' |
| ***Fdft1*** | 5'-GATCCCACTGCTGTGTAAC-3' | 5'-TTTCTAAACTCCAGGGAGATCG-3' |
| ***G6pdx*** | 5'-TACCATCTGGTGGCTGTTC-3' | 5'-TTTCGGATGTCGTCCACT-3' |
| ***Gas6*** | 5'-CCTCTGCGATGAGGGATA-3' | 5'-GACACAGGTCTGCTCAC-3' |
| ***H6pd*** | 5'-TCCAGGAGGAGGAGATGT-3' | 5'-GGTTCTGATCTCGGAAGGG-3' |
| ***Hif1a*** | 5'-TGCTTTGATGTGGATAGCG-3' | 5'-ATTCTTTGCCTCTGTGTCT-3' |
| ***Hk2*** | 5'-GAGATTGGTCTCATTGTGGGT-3' | 5'-CTCCATGTTGATGCACATCC-3' |
| ***Idi1*** | 5'-TCATCCATTAAGTAACCCAGGC-3' | 5'-CAACCTCTTCCAAGGGTAT-3' |
| ***Igf2bp2*** | 5'-GAGACCAAACTGGCTGAG-3' | 5'-TTTCTTCAGGTTTCTGCCTT-3' |
| ***Igfbp3*** | 5'-GAGTCTAAGCGGGAGACAG-3' | 5'-CAGTTTGGGATGTGGACG-3' |
| ***Igfbp4*** | 5'-CCCATTCCAAACTGTGACC-3' | 5'-CACACCAGCACTTGCCA-3' |
| ***Ikbkg*** | 5'- CCACCAAGGATCGGCAA -3' | 5'- CTGTGCTGCTGCTGTAG -3' |
| ***Ikkb*** | 5'- CAGACAGCAAGACAAACGA -3' | 5'- GACACTTTCCGGTTGGG -3' |
| ***Il-10*** | 5'-GCAGGACTTTAAGGGTTACT-3' | 5'-TCCTTGATTTCTGGGCCAT-3' |
| ***Il12b*** | 5'-AGCTGGAGAAAGACGTTTAT-3' | 5'-GTCTGAGGTCCAGGTGAT-3' |
| ***Il-1b*** | 5'-AGGCAGGCAGTATCAC-3' | 5'-CACACCAGCAGGTTATC-3' |
| ***Il1r1*** | 5'- CTGTTGGTGAGGAATGTGG-3' | 5'- TAACAGGTCTGTCCCTCTTG-3' |
| ***Il-1r2*** | 5'- GGAGACAATACCAGCATCAT -3' | 5'- TAAGCAGCCGAGATAAACG -3' |
| ***Insig1*** | 5'-GCGGCTGTTGTCGGTTTA-3' | 5'-TGATGCCAACGAACACG-3' |
| ***Irf4*** | 5'-TCCGACAGTGGTTGATCGAC-3' | 5'-CCTCACGATTGTAGTCCTGCTT-3' |
| ***Irf5*** | 5'-GGTCAACGGGGAAAAGAAACT-3' | 5'-CATCCACCCCTTCAGTGTACT-3' |
| ***Ldh1*** | 5'-ACCAATACAGAGCAACTGAT-3' | 5'-CCACCACCTTCTTCAAAGTC-3' |
| ***Ldha*** | 5'-CCTCTCTGTGGCAGACT-3' | 5'-CATTGATTCCATAGAGACCCT-3' |
| ***Ldlr*** | 5'-CAACAAGTTCAAGTGTCACAG-3' | 5'-CATTGTTGTCCAAACACTCG-3' |
| ***Mct1*** | 5'-ATGTATGCTGGAGGTCCTAT-3' | 5'-CAACCAGACAGACAACCAC-3' |

| | | |
|---|---|---|
| ***Mct4*** | 5'-GTCATCACTGGCTTGGG-3' | 5'-CATTGGCAATAGGGCGAC-3' |
| ***Mmp9*** | 5'-CTGGATAAGTTGGGTCTAGG-3' | 5'-CTTCTGAGACTTCAAGTCGAAT-3' |
| ***Mrc1*** | 5'-CTCTGTTCAGCTATTGGACGC-3' | 5'-CGGAATTTCTGGGATTCAGCTTC-3' |
| ***Mvd*** | 5'-TGCCTGAGAGAGATTCGC-3' | 5'-ATAGCTGAGGCTGAGGG-3' |
| ***Nfkb1*** | 5'- GCAACTCACAGACAGAGAGAA-3' | 5' CTGTGAACATGAGGCGCA--3' |
| ***Nfkb2*** | 5'- GCTTCAGATTTCGATATGGCT -3' | 5'- GCCGGTCCCTCATAGTT -3' |
| ***Nfkbia*** | 5'-TGGAGCACTTGGTGACT-3' | 5'- CACATTTCAACAAGAGCGAAAC-3' |
| ***Nfkbib*** | 5'- GCAGAATGACCTAGGCCAAA -3' | 5'- GCTGCATACAACTTCTCTACT -3' |
| ***Nfkbie*** | 5'- TCGAGGCGCTCACATAC -3' | 5'- GCAACAGAATAGCACCGAC -3' |
| ***Nfkbiz*** | 5'- CCGATTTCTCCTCCACT -3' | 5'- TCTCTGCTGCCTGCCAA -3' |
| ***Nlrp3*** | 5'- GACTTTGGAATCAGATTGCT -3' | 5'- GGGTCCTTCATCTTTTCAC -3' |
| ***Nos2*** | 5'-GGAATTCACAGCTCATCCG-3' | 5'-AGAGGCAGCACATCAAAG-3' |
| ***Pkm2*** | 5'-CGCCTGGACATTGACTC-3' | 5'-GCCACATTCATTCCAGACTT-3' |
| ***Rela*** | 5'- CCTTTCAATGGACCAACTG -3' | 5'- TGATGGTGCTGAGGGAT -3' |
| ***Relb*** | 5'- ATCGAGCTTCGAGACTGT -3' | 5'- ACGAGGCTATGTGGGTGTA -3' |
| ***Retnla*** | 5'-CCAATCCAGCTAACTATCCCTCC-3' | 5'-ACCCAGTAGCAGTCATCCCA-3' |
| ***Sdhb*** | 5'-CTTGTAGAGAAGGCATCTGT-3' | 5'-TTTGCTGAGGTCCGTGTC-3' |
| ***Sirt1*** | 5'-CTTACCAGAACAGTTTCATAGAGC-3' | 5'-GGGTATAGAACTTGGAATTAGTGC-3' |
| ***Socs1*** | 5'-CTGCGGCTTCTATTGGGGAC-3' | 5'-AAAAGGCAGTCGAAGGTCTCG-3' |
| ***Socs3*** | 5'-ATGGTCACCCACAGCAAGTTT-3' | 5'-TCCAGTAGAATCCGCTCTCCT-3' |
| ***Stard4*** | 5'-GATCCAGTATCACAGCATCG-3' | 5'-CACGTCATCCATAACTCCTTG-3' |
| ***Stat1*** | 5'-TCACAGTGGTTCGAGCTTCAG-3' | 5'-GCAAACGAGACATCATAGGCA-3' |
| ***Stat6*** | 5'-CTCTGTGGGGCCTAATTTCCA-3' | 5'-CATCTGAACCGACCAGGAACT-3' |
| ***Tgfb1*** | 5'-GGGAAGCAGTGCCCGAA-3' | 5'-GGTAACGCCAGGAATTGT-3' |
| ***Timp1*** | 5'-GAAAGCCTCTGTGGATATGC-3' | 5'-ATTTCCGTTCCTTAGGCG-3' |
| ***Tlr4*** | 5'- TCTTCTCCTGCCTGACAC -3' | 5'- TGGTTGAAGAAGGAATGTCATC -3' |
| ***Tnfrsf1a*** | 5'- TCCGCTTGCAAATGTCAC -3' | 5'- GGGATATCGGCACATTAAACT -3' |
| ***Tnfrsf1b*** | 5'-TACAAACCGGAACCTGGG-3' | 5'-GTCCGAGGTCTTGTTGC-3' |
