## Supplemental Figures for "The m^6^A reader IGF2BP2 directs immune-metabolic reprogramming in Leishmania amazonensis-infected macrophages"

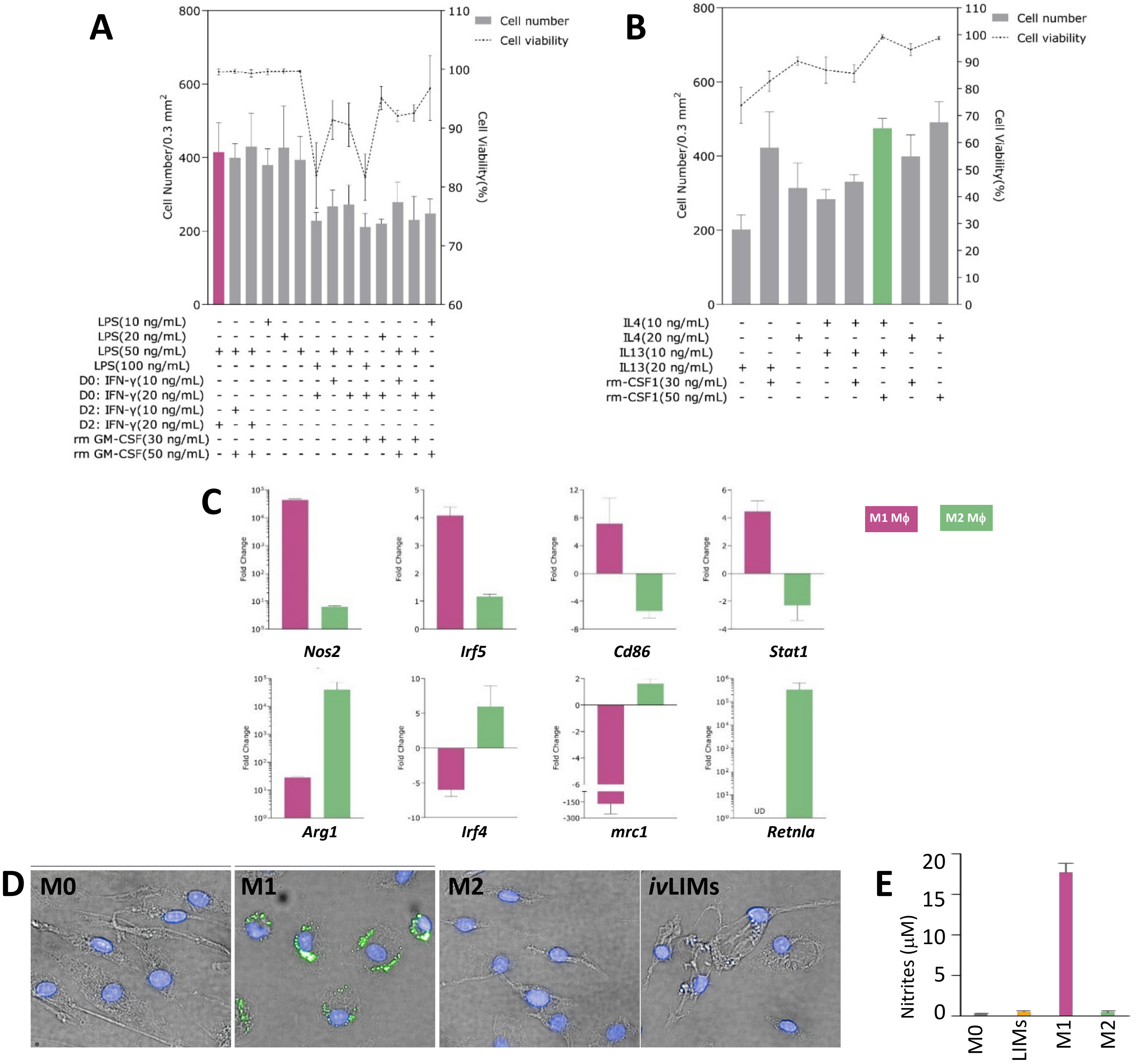


Supplementary Figure 1, Zhang S. et al.

A

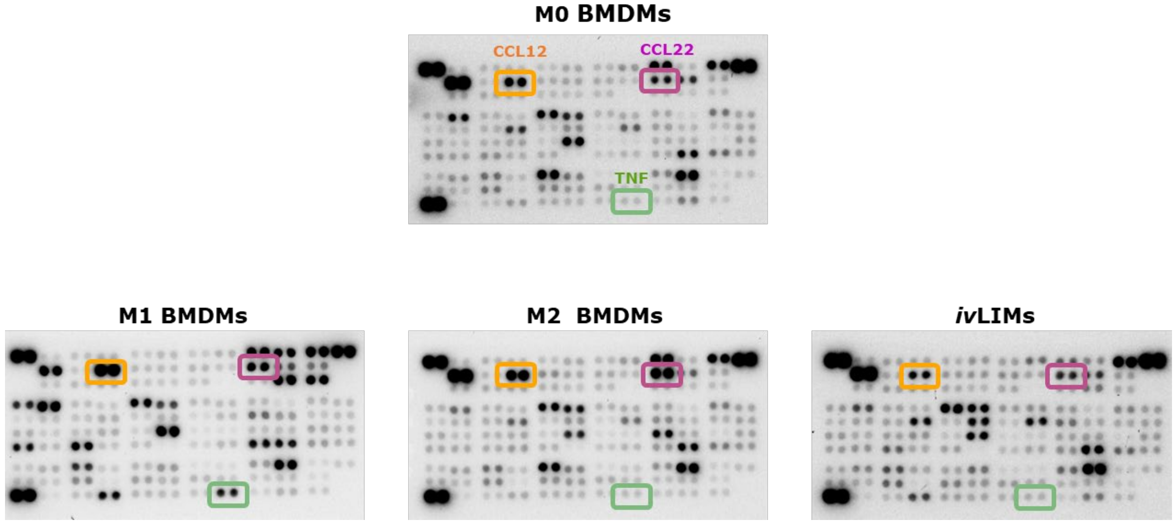


B

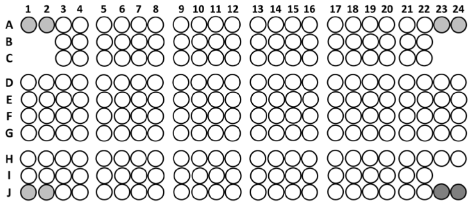


Mouse XL Cytokine Array Coordinates

C

| Coordinate | Analyte/Control | Coordinate | Analyte/Control | Coordinate | Analyte/Control | Coordinate | Analyte/Control | Coordinate | Analyte/Control |
|---|---|---|---|---|---|---|---|---|---|
| A1/A2 | Reference Spots | C1/C2 | | E1/E2 | FGF acidic | G1/G2 | IL-10 | I1/I2 | PDGF-BB |
| A3/A4 | Adiponectin/Acrp30 | C3/C4 | CD160 | E3/E4 | FGF-21 | G3/G4 | IL-11 | I3/I4 | Pentraxin 2/SAP |
| A5/A6 | Amphiregulin | C5/C6 | Chemerin | E5/E6 | Flt-3 Ligand | G5/G6 | IL-12p40 | I5/I6 | Pentraxin 3/ TSG-14 |
| A7/A8 | Angiopoietin-1 | C7/C8 | Chitinase 3-like 1 | E7/E8 | Gas6 | G7/G8 | IL-13 | I7/I8 | Periostin/OSF-2 |
| A9/A10 | Angiopoietin-2 | C9/C10 | Coagulation Factor III/Tissue Factor | E9/E10 | G-CSF | G9/G10 | IL-15 | I9/I10 | Pref-1/DLK-1/FA1 |
| A11/A12 | Angiopoietin-like 3 | C11/C12 | Complement Component C5/C5a | E11/E12 | GDF-15 | G11/G12 | IL-17A | I11/I12 | Proliferin |
| A13/A14 | BAFF/BLyS/TNFSF13B | C13/C14 | Complement Factor D | E13/E14 | GM-CSF | G13/G14 | IL-22 | I13/I14 | Proprotein Convertase 9/PCSK9 |
| A15/A16 | C1q R1/CD93 | C15/C16 | C-Reactive Protein/CRP | E15/E16 | HGF | G15/G16 | IL-23 | I15/I16 | RAGE |
| A17/A18 | CCL2/JE/MCP-1 | C17/C18 | CX3CL1/Fractalkine | E17/E18 | ICAM-1/CD54 | G17/G18 | IL-27 | I17/I18 | RBP4 |
| A19/A20 | CCL3/CCL4 MIP-1 alpha/beta | C19/C20 | CXCL1/KC | E19/E20 | IFN-gamma | G19/G20 | IL-28 | I19/I20 | Reg3G |
| A21/A22 | CCL5/RANTES | C21/C22 | CXCL2/MIP-2 | E21/E22 | IGFBP-1 | G21/G22 | IL-33 | I21/I22 | Resistin |
| A23/A24 | Reference Spots | C23/C24 | | E23/E24 | IGFBP-2 | G23/G24 | LDL R | I23/I24 | |
| B1/B2 | | D1/D2 | CXCL9/MIG | F1/F2 | IGFBP-3 | H1/H2 | Leptin | J1/J2 | Reference Spots |
| B3/B4 | CCL6/C10 | D3/D4 | CXCL10/IP-10 | F3/F4 | IGFBP-5 | H3/H4 | LIF | J3/J4 | E-Selectin/CD62E |
| B5/B6 | CCL11/Eotaxin | D5/D6 | CXCL11/I-TAC | F5/F6 | IGFBP-6 | H5/H6 | Lipocalin-2/NGAL, | J5/J6 | P-Selectin/CD62P |
| B7/B8 | CCL12/MCP-5 | D7/D8 | CXCL13/BLC/BCA-1 | F7/F8 | IL-1 alpha/IL1F1 | H7/H8 | LIX | J7/J8 | Serpin E1/PAI-1 |
| B9/B10 | CCL17/TARC | D9/D10 | CXCL16 | F9/F10 | IL-1 beta/IL-1F2 | H9/H10 | M-CSF | J9/J10 | Serpin F1/PEDF |
| B11/B12 | CCL19/MIP-3 beta | D11/D12 | Cystatin C | F11/F12 | IL-1ra/IL-1F3 | H11/H12 | MMP-2 | J11/J12 | Thrombopoietin |
| B13/B14 | CCL20/MIP-3 alpha | D13/D14 | Dkk-1 | F13/F14 | IL-2 | H13/H14 | MMP-3 | J13/J14 | TIM-1/KIM-1/HAVCR |
| B15/B16 | CCL21/6Ckine | D15/D16 | DPPIV/CD26 | F15/F16 | IL-3 | H15/H16 | MMP-9 | J15/J16 | TNF-alpha |
| B17/B18 | CCL22/MDC | D17/D18 | EGF | F17/F18 | IL-4 | H17/H18 | Myeloperoxidase | J17/J18 | VCAM-1/CD106 |
| B19/B20 | CD14 | D19/D20 | Endoglin/CD105 | F19/F20 | IL-5 | H19/H20 | Osteopontin (OPN) | J19/J20 | VEGF |
| B21/B22 | CD40/TNFRSF5 | D21/D22 | Endostatin | F21/F22 | IL-6 | H21/H22 | Osteoprotegerin/TNFRSF11B | J21/J22 | WISP-1/CCN4 |
| B23/B24 | | D23/D24 | Fetuin A/AHSG | F23/F24 | IL-7 | H23/H24 | PD-ECGF/Thymidine phosphorylase | J23/J24 | Negative control |

Mouse XL Cytokine Array list

Supplementary Figure 2, Zhang S. et al.

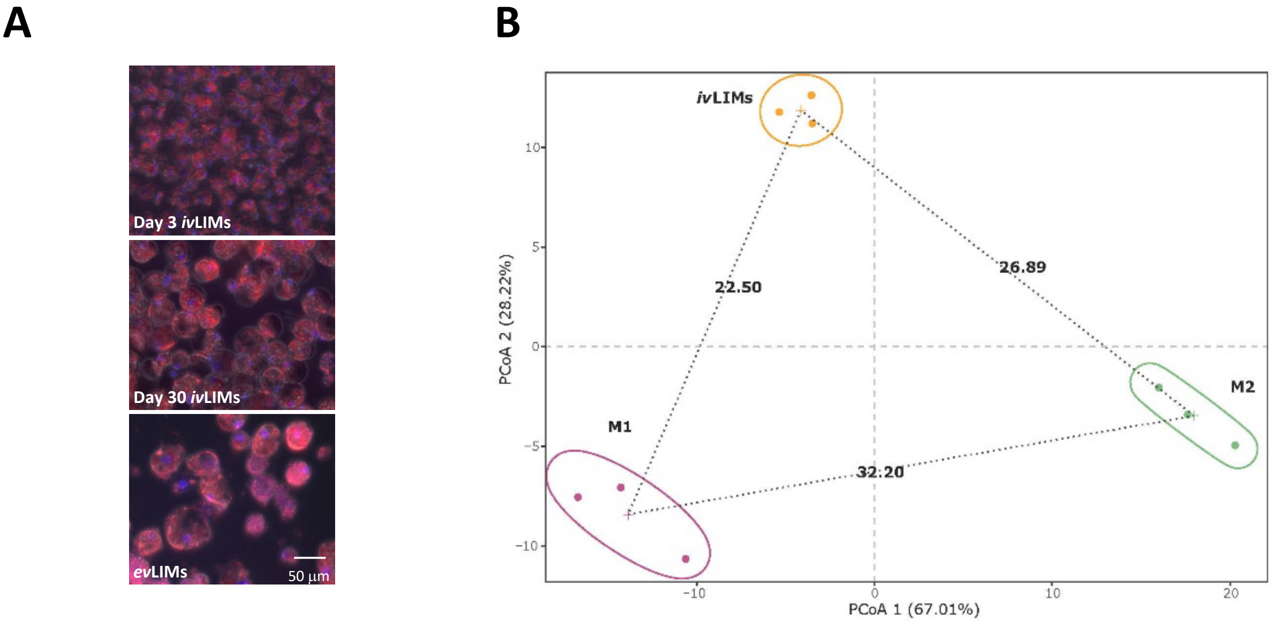


**Supplementary Figure 3, Zhang S. et al.**

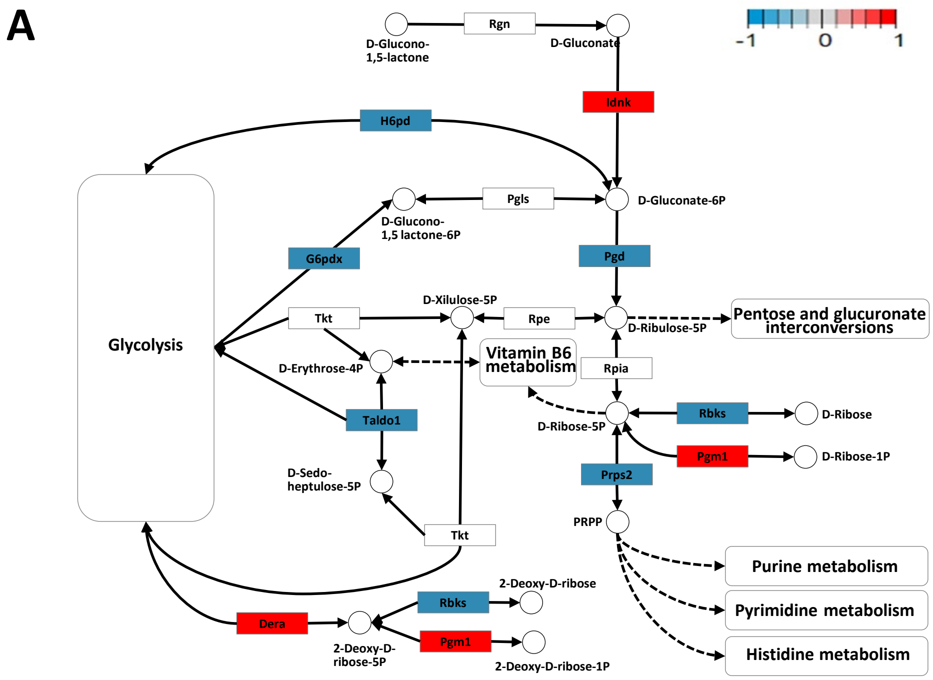


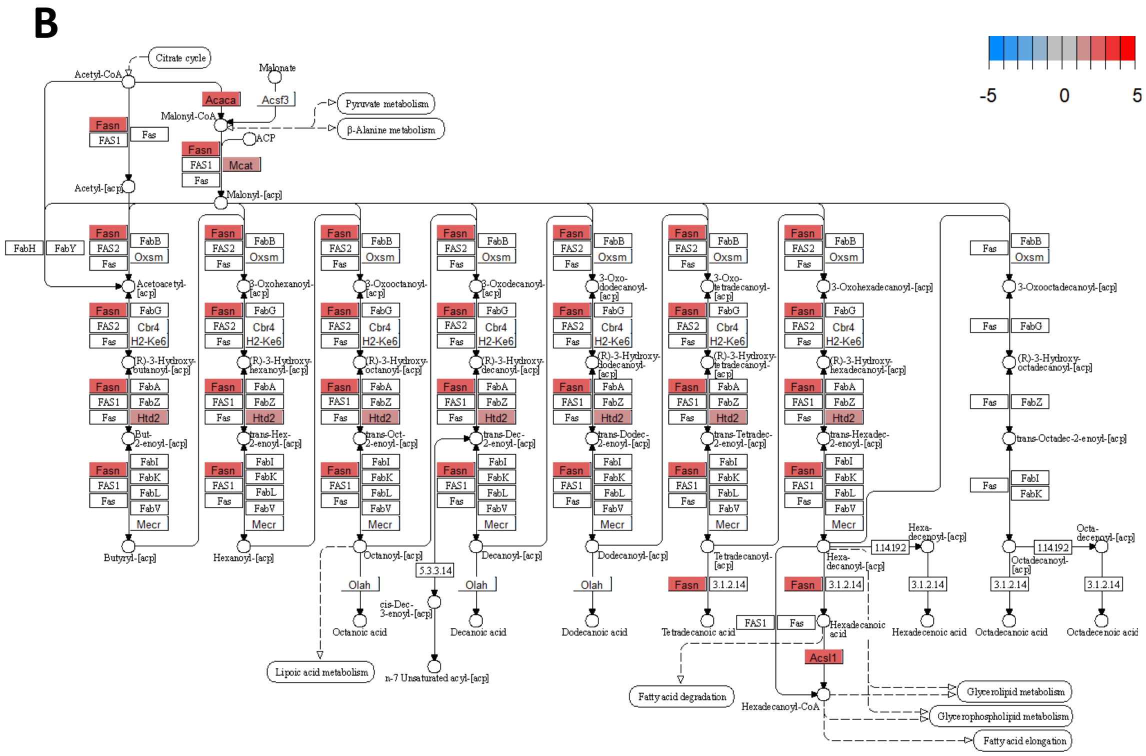


**Supplementary Figure 4, Zhang S. et al.**

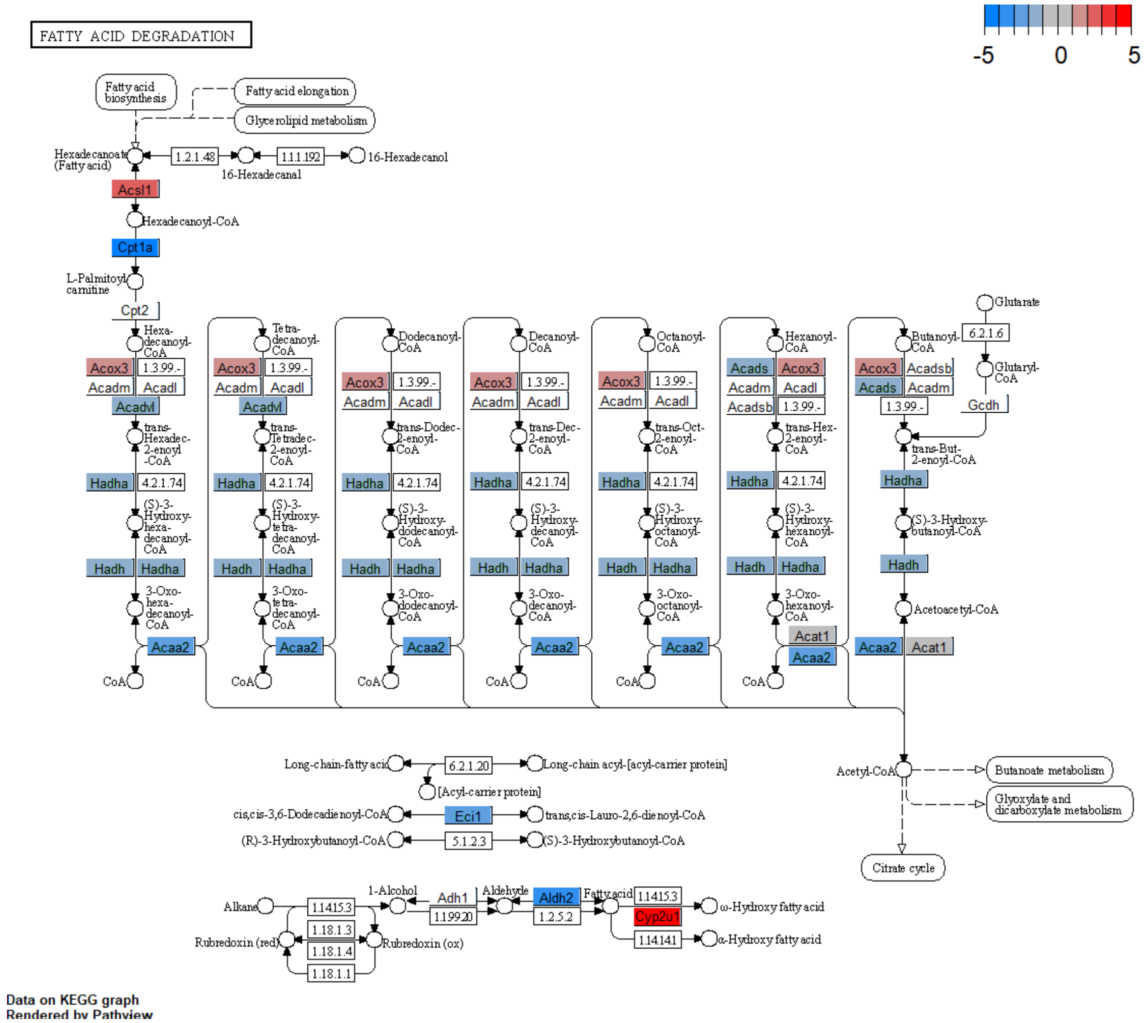


Supplementary Figure 5, Zhang S. et al.

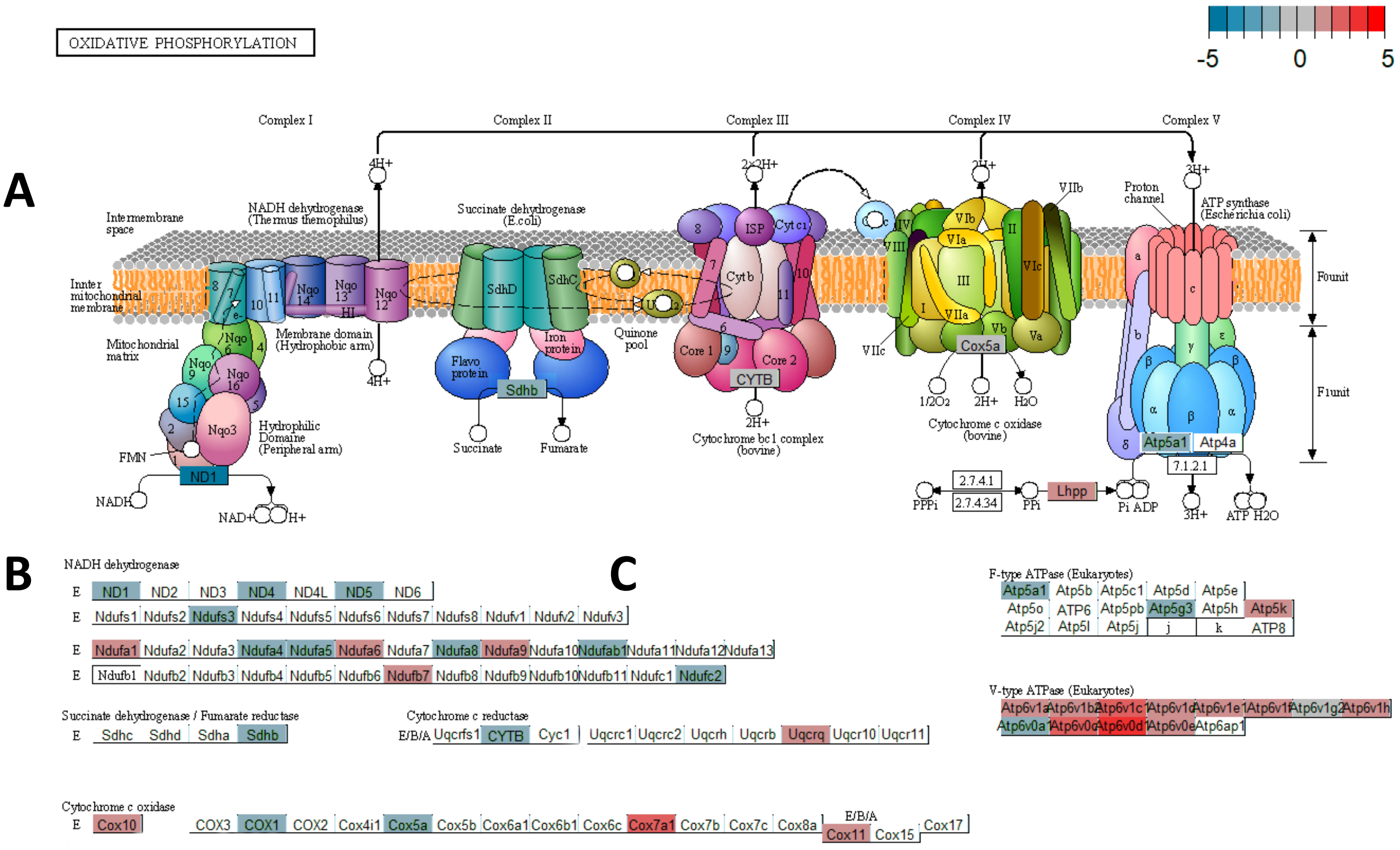


**Supplementary Figure 6, Zhang S. et al.**

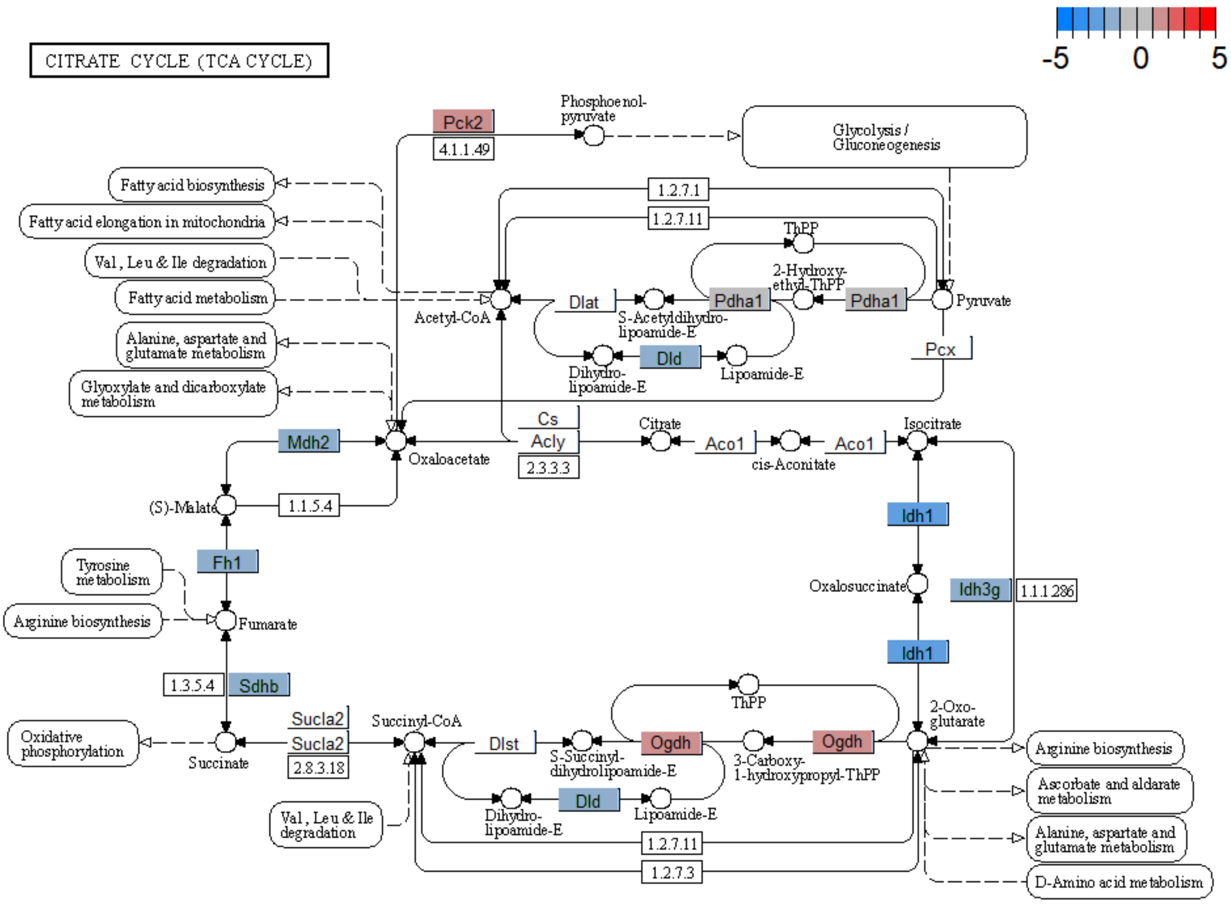


**Supplementary Figure 7, Zhang S. et al.**

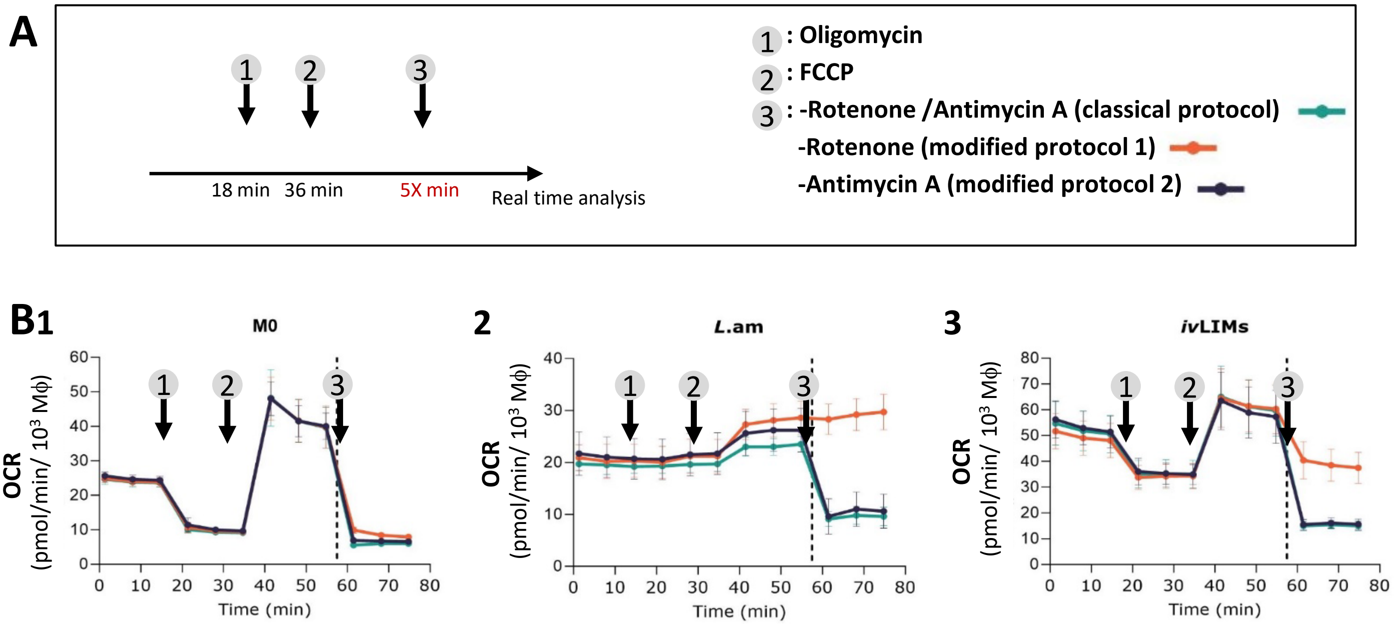


Supplementary Figure 8, Zhang S. et al.

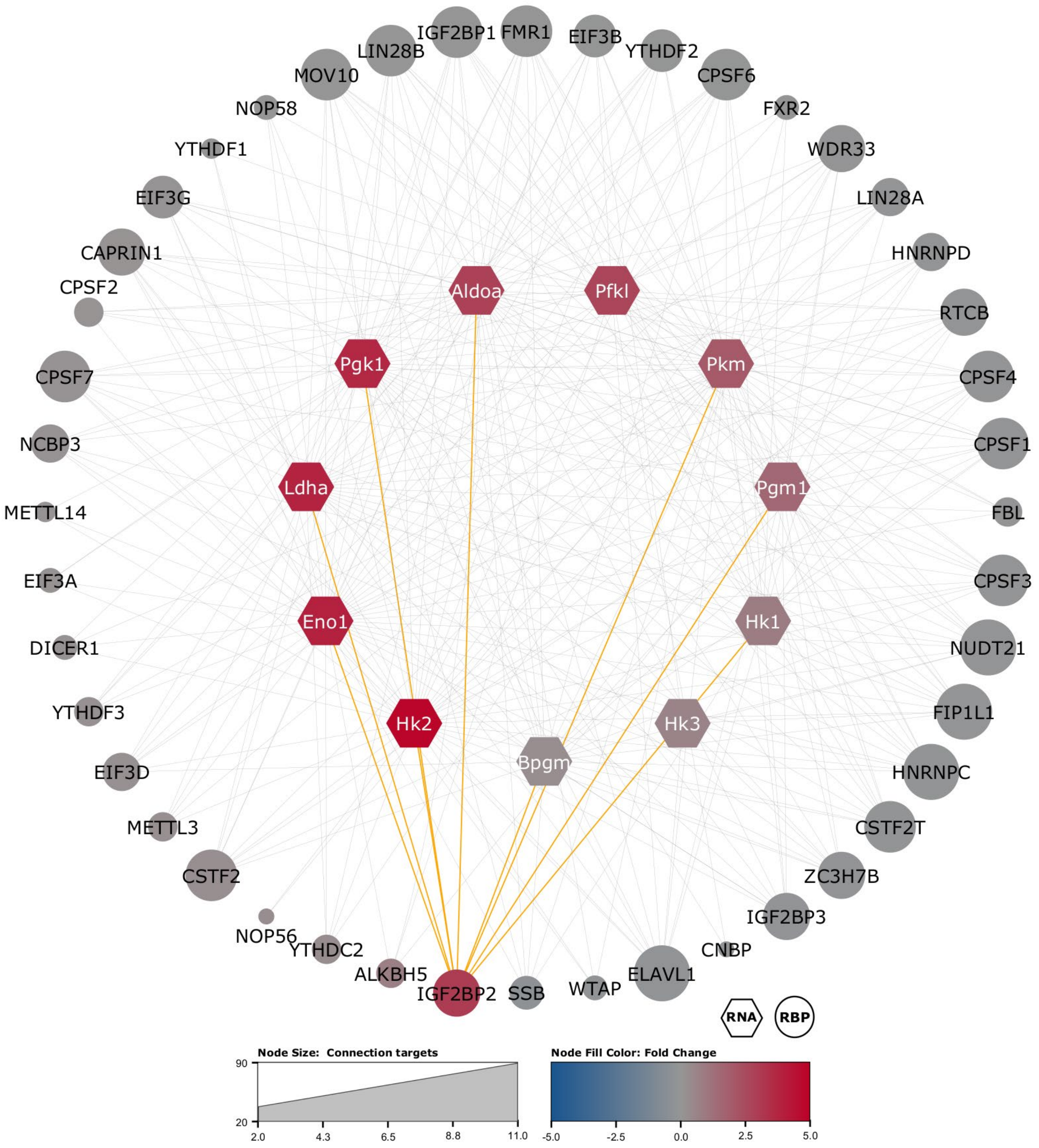


**Supplementary Figure 9, Zhang S. et al.**

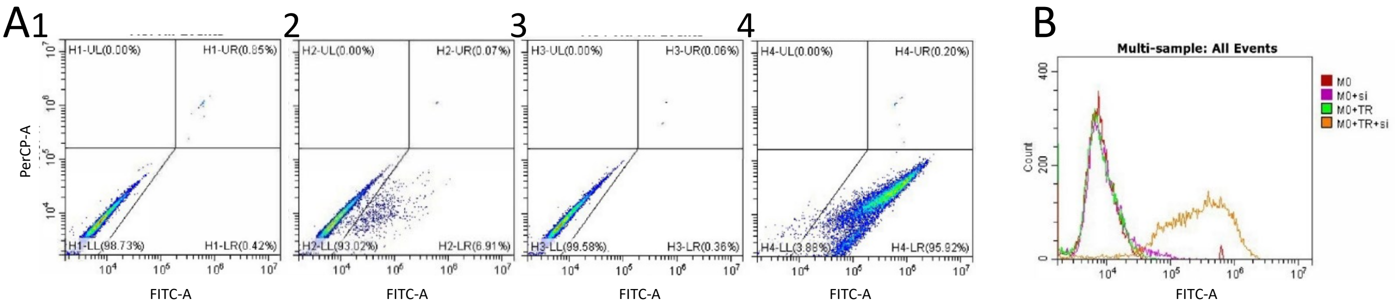


**Supplementary Figure 10, Zhang S. et al.**
